# Genome graphs detect human polymorphisms in active epigenomic state during influenza infection

**DOI:** 10.1101/2021.09.29.462206

**Authors:** Cristian Groza, Xun Chen, Alain Pacis, Marie-Michelle Simon, Albena Pramatarova, Katherine A. Aracena, Tomi Pastinen, Luis B. Barreiro, Guillaume Bourque

**Affiliations:** Quantitative Life Sciences, McGill University, Montréal, QC, Canada; Institute for the Advanced Study of Human Biology (WPI-ASHBi), Kyoto University, Kyoto, Japan; Canadian Centre for Computational Genomics, McGill University, Montréal, QC, Canada; McGill Genome Centre, Montréal, QC, Canada; Human Genetics, University of Chicago, Chicago, IL, USA; Genomic Medicine Center, Children’s Mercy Hospital and Research Institute, KC, MO, USA; Committee on Genetics, Genomics, and Systems Biology, University of Chicago, Chicago, IL, USA; Section of Genetic Medicine, Department of Medicine, University of Chicago, Chicago, IL, USA; Committee on Immunology, University of Chicago, Chicago, IL, USA; Human Genetics, McGill University, Montréal, QC, Canada

**Keywords:** epigenomics, mobile elements, pangenomics, influenza

## Abstract

Genetic variants, including mobile element insertions (MEIs), are known to impact the epigenome. We hypothesized that the use of a genome graph, which encapsulates genetic diversity, could reveal missing epigenomic signal. Given the contributions of mobile elements to the evolution of primate innate immunity, we tested this in monocyte-derived macrophages obtained from 35 individuals before and after *Influenza* virus infection. After characterizing genetic variants in this cohort using linked-reads, including 5140 Alu, 316 L1, 94 SVAs and 48 ERVs, we incorporated them into a genome graph. Mapping epigenetic data to this graph revealed 2.5%, 3.0% and 2.3% novel peaks for H3K4me1 and H3K27ac ChIP-seq and ATAC-seq respectively. Notably, using a genome graph also modified quantitative trait loci estimates and we observed 375 polymorphic MEIs in active epigenomic state. For example, we found an AluYh3 polymorphism whose chromatin state changed after infection and that was associated with the expression of *TRIM25*, a gene that restricts influenza RNA synthesis. Our results demonstrate that graph genomes can reveal regulatory regions that would have been overlooked by other approaches.

## Introduction

Structural variants (SVs) contribute the largest number of variable nucleotides in an individual [1], have larger effect sizes on gene expression [2] and are associated with functionally relevant epigenetic differences between humans and chimpanzees [3]. A particular class of SVs, mobile element insertions (MEIs), likely influences the epigenome since fixed mobile elements are known to harbor transcription factor binding sites [4, 5] and have contributed primate specific regulatory regions [6]. The epigenetic features that occur on SVs are not immediately accessible when mapping to a linear and incomplete reference genome [7, 8] but could potentially be accessed using a graph genome. Indeed, using the personal genome of a single individual we have shown previously that genome graphs can recover epigenomic signal in genetically variable regions of the genome [9]. Genome graph approaches have also been used to find differential CpG methylation within SVs in twelve medaka fish genomes [10].

Obtaining accurate maps of SVs with short read libraries can be challenging for several reasons [11]. First, repeats are abundant in eukaryotic genomes and resolving variation in these regions can be more difficult since read mapping is often ambiguous. Second, mapping short reads reveals only the break points of insertions and does not provide their actual sequence without assembling the reads into larger contigs. Third, short read assembly algorithms cannot distinguish between highly similar sequences and tend to collapse copy number variation. To mitigate these shortcomings, paired-end and linked read libraries [12] have been developed. Linked read libraries go further than paired-end libraries by labeling each read with a barcode that represent the DNA fragments from which it originates. This provides long range positional information in regions of the human genome that cannot be reached by short reads alone. Linked reads have been used to genotype [13, 14, 15], identify SVs [16, 17, 18], detect MEI polymorphisms [19] and assemble genomes [20, 21].

Given that mobile elements have been found to be co-opted in innate immunity [22], we decided to study the impact on the epigenome of genetic variants and MEIs in the response to *Influenza* virus (IAV) infection. Specifically, we used data obtained from monocyte-derived macrophages from 35 individuals of African- or European-descent before and after *in-vitro* IAV infection [23]. This included whole-genome sequence (WGS) data together with H3K4me1, H3K27ac ChIP-seq, ATAC-seq and RNA-seq data to characterize the transcriptome and the chromatin state. First, we developed a new approach to build a genome graph that includes MEIs by resolving the sequence of insertions using locally assembled linked reads (Fig S1). We validated the method by generating ChIP-seq and ATAC-seq data from the NA12878 benchmark genome [24] following the protocols used in Aracena et al. [23]. Next, we generated linked read data for the 35 individuals in the IAV infected cohort and applied our new approach to build a graph that includes SNPs, indels and MEIs. Using this genome graph, we showed we could identify regulatory sequences that would have been missed otherwise.

## Results

### Adding MEIs to the NA12878 genome graph reveals additional epigenomic signal

We chose the NA12878 genome to develop and benchmark our approach since WGS, linked reads and a haplotype resolved assembly were already available [24]. First, we ran MELT and ERVcaller on paired-end WGS data to identify and genotype 2175 Alu, 351 LINE1, 106 SVA and 6 ERV insertions (Fig 1A). Of these calls, 66% (1738) were previously observed by Ebert et al. [25]. MELT and ERVcaller could only predict the breakpoints of MEIs and to obtain the sequence of these insertions, we developed a local linked read assembly tool, which we called `BarcodeAsm` (Methods). By applying this tool, we successfully assembled the sequences of 1054 Alu, 117 LINE1, 17 SVA and 35 ERV instances (Fig 1B), sometimes recovering both copies of homozygous insertions corresponding to 963 loci (Fig 1A, right). We noted an excess of ERV annotations due to a set of misannotated short sequences that appear with low frequency in the local assembly windows. Next, to validate these assembled insertions, we aligned them against the haplotype resolved assembly of NA12878 and calculated the proportion of their length that matched (Fig 1C). Despite expecting some errors in our local assembly and in the *de novo* assembly, we found that 925 loci (96.1%) had a match over 95% of their length. These matches were also equally distributed between the two haplotypes of the *de novo* assembly, confirming the robustness of the approach.

**Figure 1:**
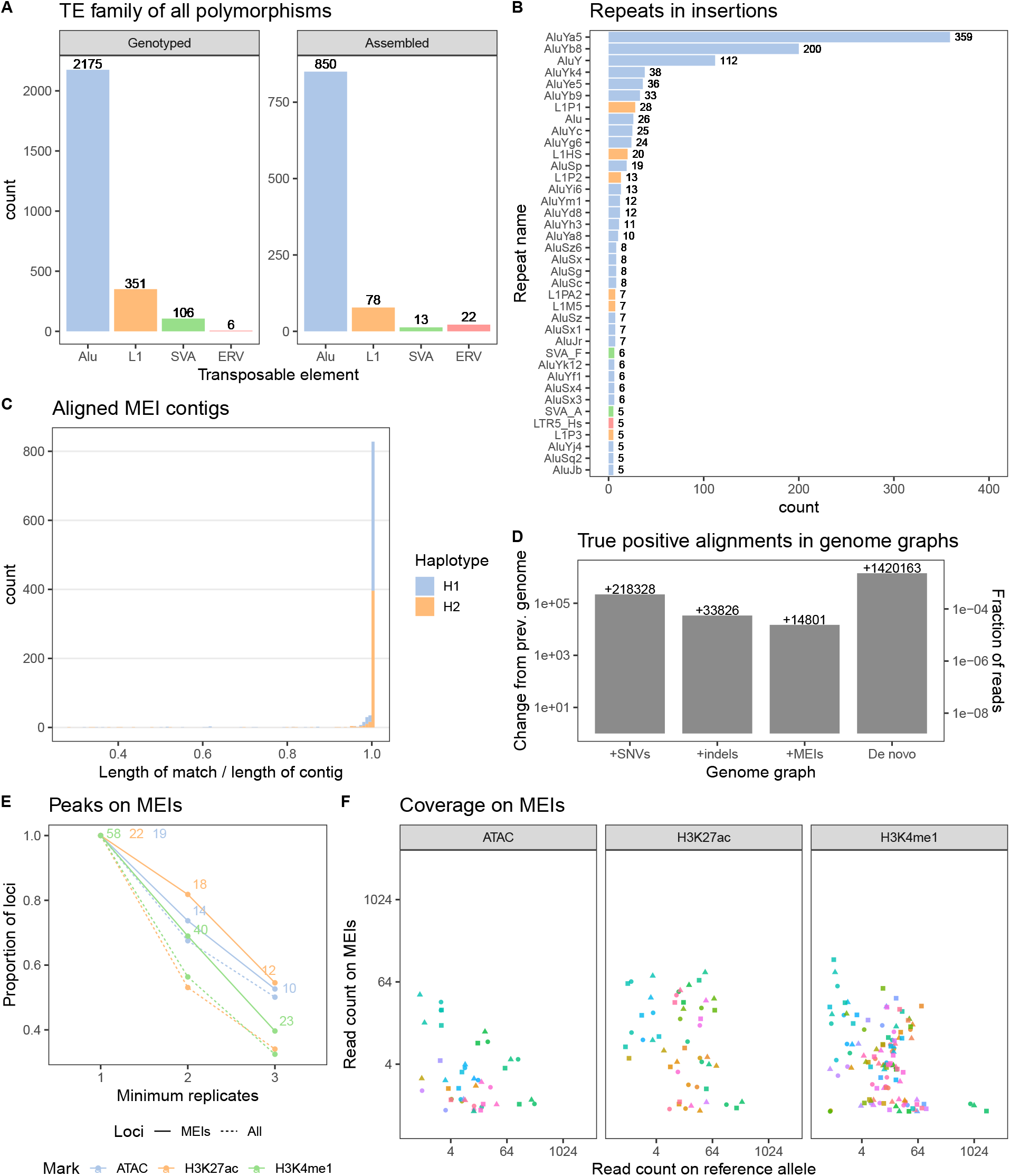
Adding MEIs to the NA12878 genome graph reveals additional epigenomic signal. A) The number of MEI breakpoints detected using ERVcaller and MELT and the full-length MEIs recovered by BarcodeAsm. B) The reassembled families ordered by frequency in the NA12878 genome. Homozygous insertions that are assembled twice are double counted. C) The spans of MEI contigs that could be matched and confirmed in the haplotype resolved assembly of NA12878. D) Change in the number of true positive alignments relative to the previous genome in increasingly complete genomes of NA12878, starting with the reference as the baseline. E) Proportion and number of peaks on MEI (full line) that were called at least once, twice and three times in the replicates. Proportion also shown for all peaks (dashed lined). F) MEI and reference allele coverage at the peak calls that overlap MEIs, stratified by locus (color) and replicate (shape).

Previously, we introduced an axis that orders reference genomes according to how similar they are to the truth [9]. This axis ranged from the least accurate sequence (the reference genome) to the most representative (the *de novo* assembly). As intermediate stages along this axis, we now defined multiple genome graphs: a +SNVs graph containing only SNVs, a +indels graph containing SNVs and indels, and a +MEIs graph containing SNVs, indels and MEIs (Methods). We also created a *de novo* graph with all the variants called in the NA12878 haplotype-resolved assembly. Next, we simulated 600 million standard WGS reads from this *de novo* graph and aligned them to the genomes along this axis. As expected, we observed that the number of true positive alignments increased as the graphs became more complete (Fig 1D). The +SNVs graph (with 3.5 × 10^6^ SNVs) correctly finds around 2.2 × 10^5^ more true alignments (+0.062 per SNV) compared to the reference graph. The +indels graph (with 5.2 × 10^5^ indels), adds 3.3 × 10^4^ true alignments (+0.063 per indel) over SNVs alone. The impact of SNVs and indels is similar because most indels are short and allelic bias in indels overcomes SNVs only at longer lengths [26]. The +MEIs graph (with 963 MEIs) adds another 1.4 × 10^4^ true alignments (+14.5 per MEI) on top of SNVs and indels, a much larger impact per variant as compared to SNVs and short indels. Finally, the *de novo* graph outperforms our best genome by 1.4 × 10^6^ true alignments since it represents even more SVs.

Having established that our genome graph recovers more true mappings, we looked for MEIs that support active histone marks and chromatin accessibility signal. To mimic the data obtained from [23], we used cells derived from NA12878 and generated three replicates for H3K4me1, H3K27ac ChIP-seq and ATAC-seq (Methods). These data were then mapped to the MEI graph and we called peaks using Graph Peak Caller [27]. We observed 58 H3K4me1, 22 H3K27ac and 19 ATAC-seq peaks that overlapped MEIs in at least one of the replicates (Fig 1E). Similarly to other peaks, roughly half were observed in all three replicates. Most loci were covered on both the MEI and the reference allele and a smaller subset were covered on only one of the alleles (Fig 1F). While the number of events is small in a single genome, they show that using a graph we can profile the chromatin in regions that are missing from the reference.

### Linked reads recover the sequences of polymorphic MEIs in a cohort

We looked to extend the method and apply it to a cohort with the 35 individuals that were exposed to IAV infection [23]. First, using MELT and ERVcaller on the WGS data, we identified and genotyped 7362 Alu, 1344 LINE1, 649 SVA and 19 ERV insertion loci (Fig 2A). Next, we generated linked read data from the same individuals and introduced a population consensus approach before attempting to identify the final MEI sequence at each locus with `BarcodeAsm` (Methods). Using this approach we were able to assemble and annotate the insertions for 5140 Alu, 316 LINE1, 94 SVA and 48 ERV (Fig 2A, right). The population consensus approach allowed us to recover a larger fraction (60% versus 37%) of MEIs because the number of attempts to reassemble a locus is equal to the frequency of the MEI allele in the cohort. Consequently, while singleton MEIs are the most numerous, they are also the least likely to be assembled (Fig S2). As expected, the length distributions of the consensus insertions identified include the Alu peak at 300 bp and a long tail associated with longer truncated and full length MEIs such as the LINE1 (Fig 2B-C). The resulting multiple sequence alignments showed few ambiguous nucleotides, suggesting that the consensus insertions are representative of most samples (Fig S3A). When we ordered mobile element families based on abundance (Fig 2D), we observed unsurprisingly that AluY families were the most common amongst Alus and that the L1HS sub-family was ranked highly [28, 29]. Similarly, the human specific SVA F sub-family [30] was found to be the most abundant among SVA families in our dataset.

**Figure 2:**
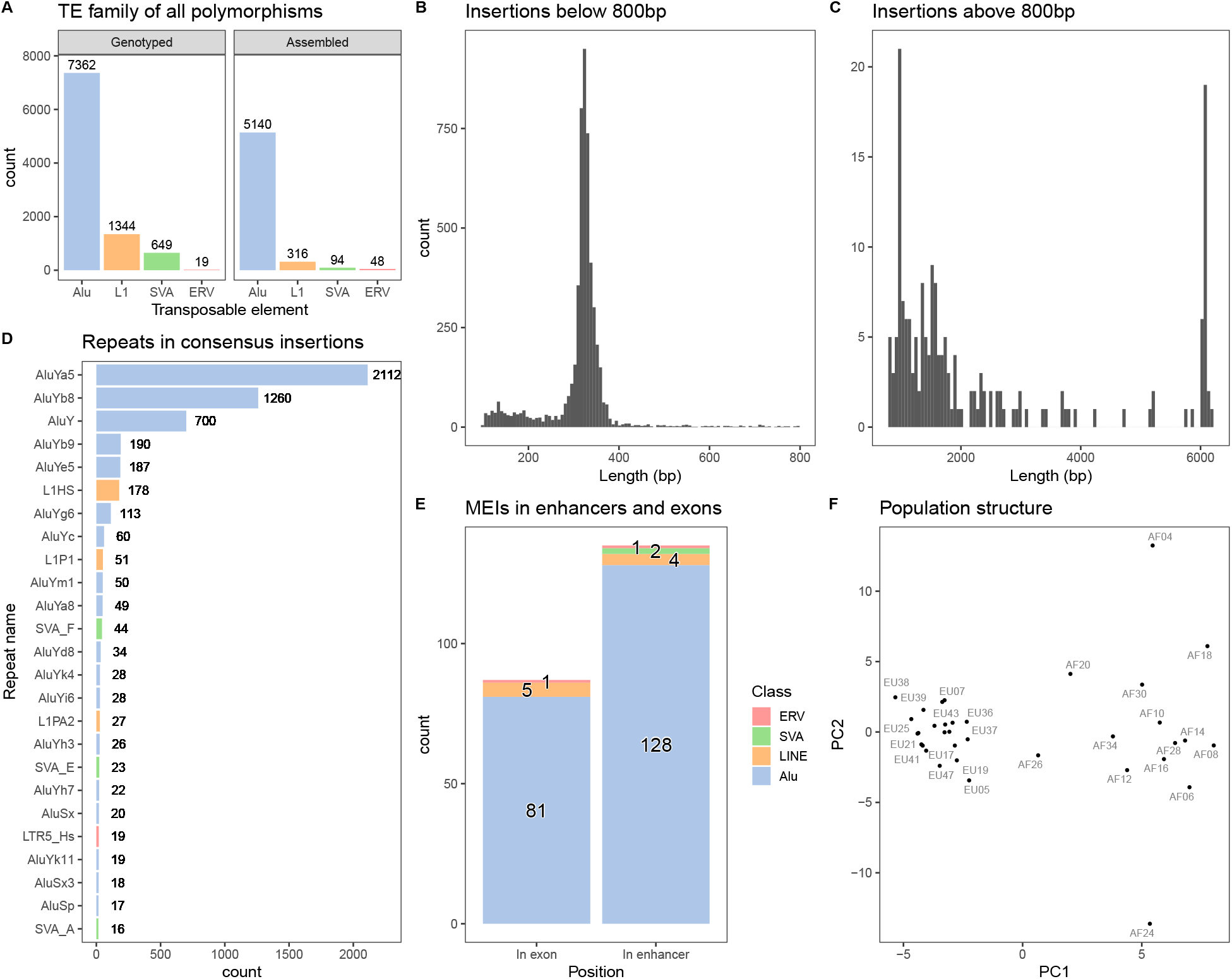
BarcodeAsm recovers the sequences of polymorphic MEIs in a cohort. A) The number and family of MEIs genotyped from short read sequencing data using ERVcaller and MELT in the entire cohort. The lengths of assembled MEIs B) below 800 bp and C) above 800 bp. D) The reassembled families ordered by frequency in the cohort. E) Number of MEIs inserted in enhancers or exons. F) The observed population structure in MEI genotypes as projected by principal component analysis.

Next, we looked at the distance between insertions and the nearest exon, enhancer and repetitive sequences in the reference genome (Methods). We found that 87 insertions, mostly Alus, were within the bodies of exons (Fig 2E), which is notable since Alus are associated with alternative transcription events [31]. We also observed 135 insertions located within enhancers (Fig 2E) and that more than half of the insertions (2941) were nested within a repetitive sequence (data not shown). Overall, the distribution of these MEIs around exons, enhancers and other repeats is similar to those in the 1000 Genomes Project [32] (Fig S3B-D). Lastly, we checked if the insertions that we reconstructed with `BarcodeAsm` and their genotype would recapitulate the known African and European ancestry structure of the cohort. We found two clusters in the principal component analysis (Fig 2F) that were consistent with the genetic ancestry and what we see from SNV calls from WGS data (Fig S4). Overall this confirms that we were able to recover a high quality set of MEIs for our cohort together with their assembled sequences.

### A genome graph increases the number of peak calls near immune genes and impacts QTL estimates

We wanted to explore the extent to which a cohort genome graph would impact read mapping and peak calling for epigenomic datasets. As expected [32], we observed more variants in samples from African ancestry (Fig 3A). Combining all these variants, we built a genome graph containing a total of 1.6 × 10^7^ SNVs, 1.1 × 10^6^ insertions, 1.3 × 10^6^ deletions and 5.6 × 10^3^ MEIs. We then mapped ChIP-seq and ATAC-seq datasets before and after IAV infection [23] and observed a decrease in unmapped reads and an increase in perfectly mapped reads that can reach up to 0.15% of reads in a data set (Fig 3B). Next, we called peaks using either the reference genome or the cohort genome graph. Peaks called with both approaches were called `common peaks`, while those called only in the reference graph are `ref-only` and those called only in the cohort graph are `graph-only`. Among H3K4me1 samples, we observed an average of 4700 (2.5%) graph-only peaks and 2200 (1.2%) ref-only peaks per sample (Fig 3C, S5A). The net increase in the number of peaks is caused by additional mapped reads that push peaks above the significance threshold. The fact that graph-only peaks are approximately twice as numerous as ref-only is consistent with what was observed previously [9]. For H3K27ac, we counted 1800 (3.0%) graph-only peaks and 1100 (1.9%) ref-only peaks (Fig S6A, S5B) on average. Among ATAC-seq datasets, graph-only events averaged 4000 (2.3%) peaks in flu-infected samples and ref-only events average 2400 (1.4%) peaks per sample (Fig S6B). In non-infected samples, the same numbers and proportions are roughly halved (S5C). We suspect that this is linked to cell death in flu-infected samples introducing cell-free DNA and more background in the ATAC-seq library preparation. Consistent with this hypothesis, infected samples show an excess of low quality peaks relative to non-infected samples (Fig S6E-F).

**Figure 3:**
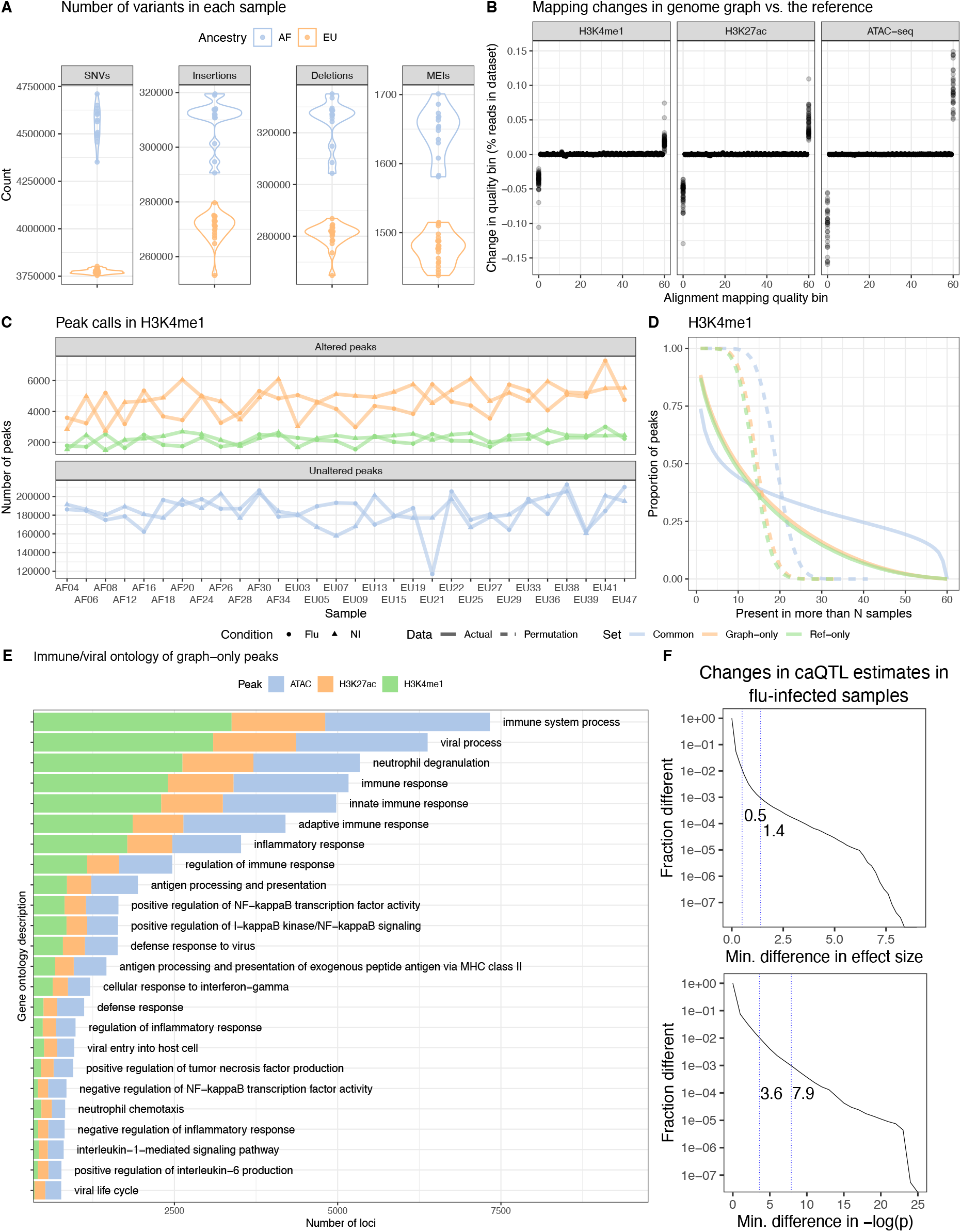
Genome graphs with millions of variants impact downstream results. A) Number of SNVs, insertions, deletions and MEIs in the cohort graph for each sample. B) Changes in the distribution of read mapping quality when using a cohort genome graph relative to the reference genome. Each point is an individual sample. C) The number of H3K4me1 altered (graph-only, ref-only) and unaltered (common) peaks between the cohort and the reference genome graphs, stratified between flu-infected and non-infected (NI) read sets. D) Inverse cumulative distributions describing how many peaks are observed in more than a number of samples. Curves that are expected by chance are also shown (dashed lines). E) Immune related gene ontology descriptions of genes within 10 Kbp of graph-only peaks. One gene may contribute multiple descriptions. F) Distributions showing caQTL effect size and p-value estimates that changed by a minimum amount between the genome graph and the reference. First and second vertical lines mark the 99th and 99.9th percentiles.

To confirm whether altered peaks were associated with sequence variants, we estimated the influence of genotype on common and graph-only peaks by logistic regression on SNPs and indels within the peaks while controlling for peak width (Table S1). We found that the log-odds for a peak to be graph-only increased by 0.11 to 0.19 when SNPs were present and by 0.45 to 0.56 when indels were present. Overall this confirms that indels have a stronger influence on peak calling as compared to SNPs. A similar analysis could not be performed with MEIs because of the small numbers. This multi-sample epigenomic dataset was also an opportunity to better understand how reliable the altered peaks were compared to common peaks. We contrasted the population frequency of graph-only, ref-only and common peaks with a peak replication cumulative distribution curve (Methods). H3K4me1, H3K27ac and ATAC peak sets generated very similar curves, with common peaks having the longest tails followed by graph-only and ref-only peaks, with the curves expected by chance decaying the fastest (Fig 3D, S6C-D). Under the random simulations, none of the peaks were observed in more than 20 datasets, but a proportion of graph-only peaks were replicated in more than 40 datasets (each individual has a non-infected and an infected dataset).

To understand the relevance of the newly identified peaks in the response to IAV, we retrieved the ontological descriptions of genes within 10 Kbp of a peak (Methods). As expected, we found that the genes near common peaks were functionally enriched for immune biological processes (Fig S7). Notably, genes near graph-only peaks were enriched for similar ontological terms (Fig S7 and Fig 3E). For example, 281, 203 and 249 genes related to “positive regulation of NF-kappaB” had graph-only peaks for H3K4me1, H3K27ac and ATAC, respectively. Finally, the use of a genome graph could affect methods to identify quantitative trait loci (QTLs), which aim to characterize the impact of genetic variants [33]. To measure this, we mapped caQTLs (chromatin accessibility) and hQTLs (H3K4me1 and H3K27ac histone modifications) using reference-based and graph-based read count estimates (Methods). We measured changes in effect size and p-value and found that while most QTLs remained the same, some do change. For example, when mapping caQTLs, the estimated effect size changes by 1.4 or more for one QTL in 1000 and the observed p-value (as −*log*(*p*)) changes by 7.9 or more (Fig 3F). The mapping of H3K4me1-QTLs and H3K27ac-QTLs is similarly affected (Fig S8). We checked the genes that are nearby these top changing QTLs and found enrichment for mostly immune pathways, which is consistent with the cell type (Fig S9). For example, the gene *HLA-DQA1* is near such changing QTLs for H3K4me1, H3K27ac and ATAC-seq. Therefore, removing reference bias from the analysis of epigenomic data using genome graphs can reveal novel peaks and improve QTL discovery.

### Genome graphs measure epigenomic signal on MEIs

We wanted to focus next on the MEIs that were assembled and introduced in the cohort genome graph. First, we took advantage of the genotypes of the individuals in the cohort to see how specific read mapping was in these repetitive and polymorphic sequences. We did this by aligning whole genome sequencing reads to the genome graph and regenotyping the MEIs in each sample (Methods). Reassuringly, we find that the graph genotypes are highly consistent with the calls made by MELT and ERVcaller (Fig 4A). When genotyping, the graph recapitulates between 1274 to 1563 genotypes per sample and only misses 60 to 95 insertions. The graph also gains between 110 and 196 insertions per sample. These gained genotypes are not necessarily false positives and could be explained by an increase in sensitivity since the exact location and sequence of these polymorphisms were known *a priori* with the graph genotyping algorithm but needed to be found *de novo* by MELT and ERVcaller. Having confirmed that read mapping was reliable in MEIs, we extracted H3K4me1, H3K27ac ChIP-seq and ATAC-seq peaks overlapping MEIs (Fig 4B, S10A-B). The peaks were labeled either as reference peaks, biallelic peaks or MEI peaks (binomial test, Methods). Notably, we found 714 MEI peaks for H3K4me1, 191 for H3K27ac and 316 for ATAC-seq. As an additional negative control, we repeated the same with peaks in MEI loci for which the samples were homozygous reference. We found that reads in these peaks were overwhelmingly assigned to the reference allele (Fig S10C-E).

**Figure 4:**
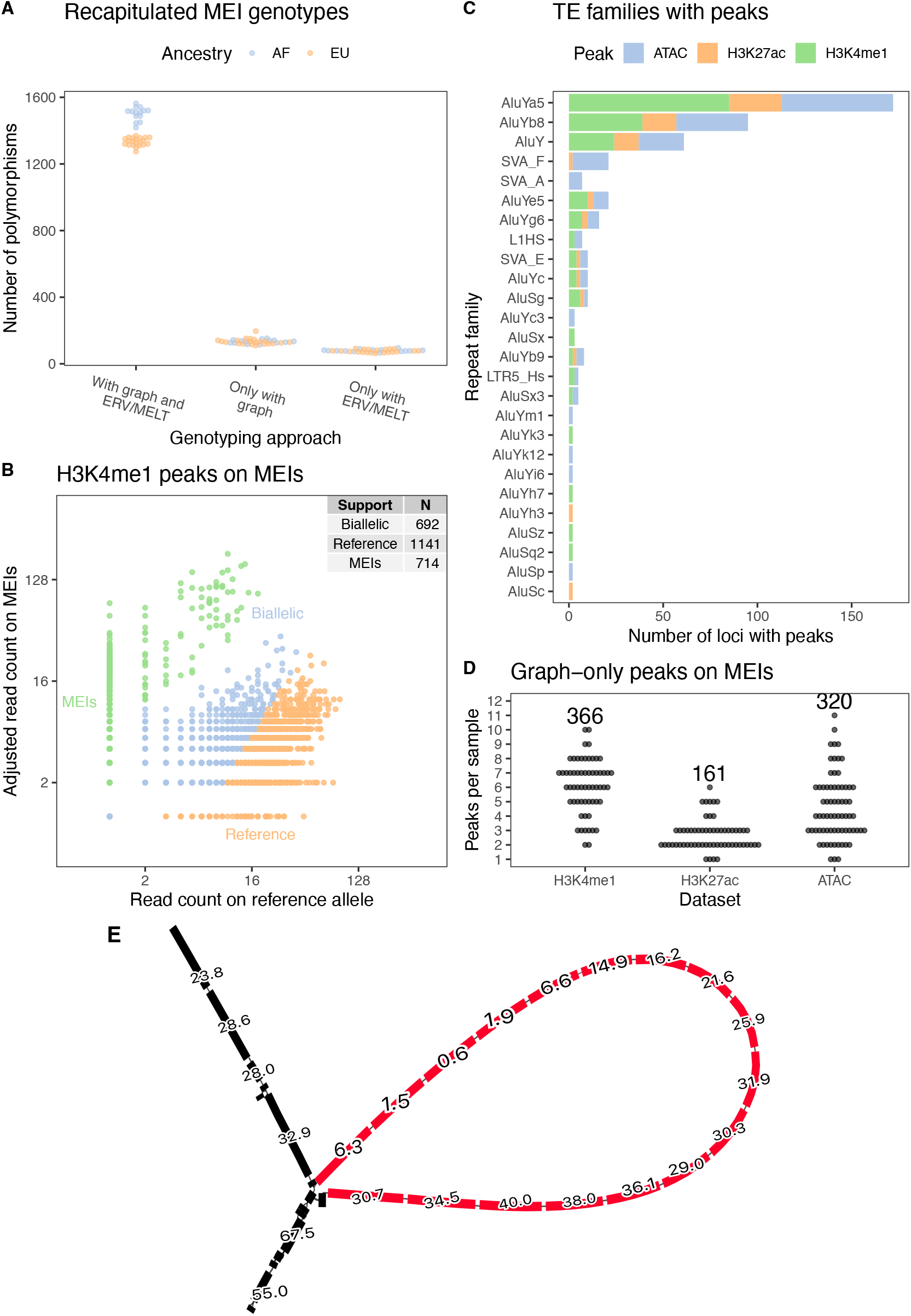
Genome graphs measure epigenomic signal on MEIs. A) Assembled MEIs that were regenotyped using the cohort genome graph. Graph genotypes are compared to the previous genotypes that were called with ERVcaller and MELT. B) Partitioning of reads between the reference and alternative allele in peaks that overlap heterozygous or homozygous MEIs. C) TE families that support peaks in at least one sample. Singletons not shown. D) Number of graph-only peaks that overlap MEIs in each sample, with total number across genomes annotated at the top. E) A graph region that represents a MEI locus. The black nodes represent the reference sequence and the red nodes represent the Alu insertion carried by some samples. The numbers show the average number of H3K4me1 reads that were mapped to each nucleotide over the span of the node. Nodes are at most 32 bp long.

MEIs peaks for H3K4me1 and H3K27ac were mostly from AluY families or other Alu elements (Fig 4C). L1 (L1HS, L1PA2) and ERV (LTR5 Hs) insertions also support a small number of peaks. Furthermore, we detected 30 SVA insertions in open chromatin states (Fig 4C). Notably, Chen et al. [34] found that SVA families become enriched in open chromatin after influenza infection in macrophages and are variable between individuals (Fig S11). Consistent with this, the overwhelming majority of ATAC-seq peaks that lie on SVA insertions were detected in the infected condition (92 of 108 peaks, 85.2%). Finally, we counted how many of these peaks were detected with the graph genome and would have been missed by a traditional approach. In total, we tallied 366 H3K4me1, 161 H3k27ac, and 320 ATAC graph-only peaks (Fig 4D) on MEIs. All together, 22.0% of H3K4me1 MEI peaks, 44.5% of H3K27ac MEI peaks and 44.0% of ATAC MEI peaks were graph-only. On the other hand, MEIs rarely disrupt peaks in the reference graph (Fig S10F). We show one Alu insertion in the graph that supports a H3K4me1 graph-only peak (Fig 4E), and its linear surjection (Fig S12).

### Cohort data reveals MEIs that act as potential enhancers

So far, we have focused on detecting single MEI alleles that support chromatin marks in individual genomes. Next, we wanted to obtain an overview of MEIs at the level of the entire cohort. In total, we identified 375 MEIs that carry at least one epigenomic peak. Specifically, 218 MEIs support H3K4me1 peaks, 90 support H3K27ac peaks and 210 support ATAC peaks (Fig 5A), many of which were found in more than one sample (Fig S13). Among the H3K4me1 MEI peaks, 58 were unique to flu-infected samples, 41 were unique to non-infected samples and 119 were found in both conditions (Fig 5B). Peaks in H3K27ac and ATAC-seq were also well balanced between the two conditions. Ordering MEIs by allele frequency, shows variable levels of occupancy by peaks (Fig 5A, right side).

**Figure 5:**
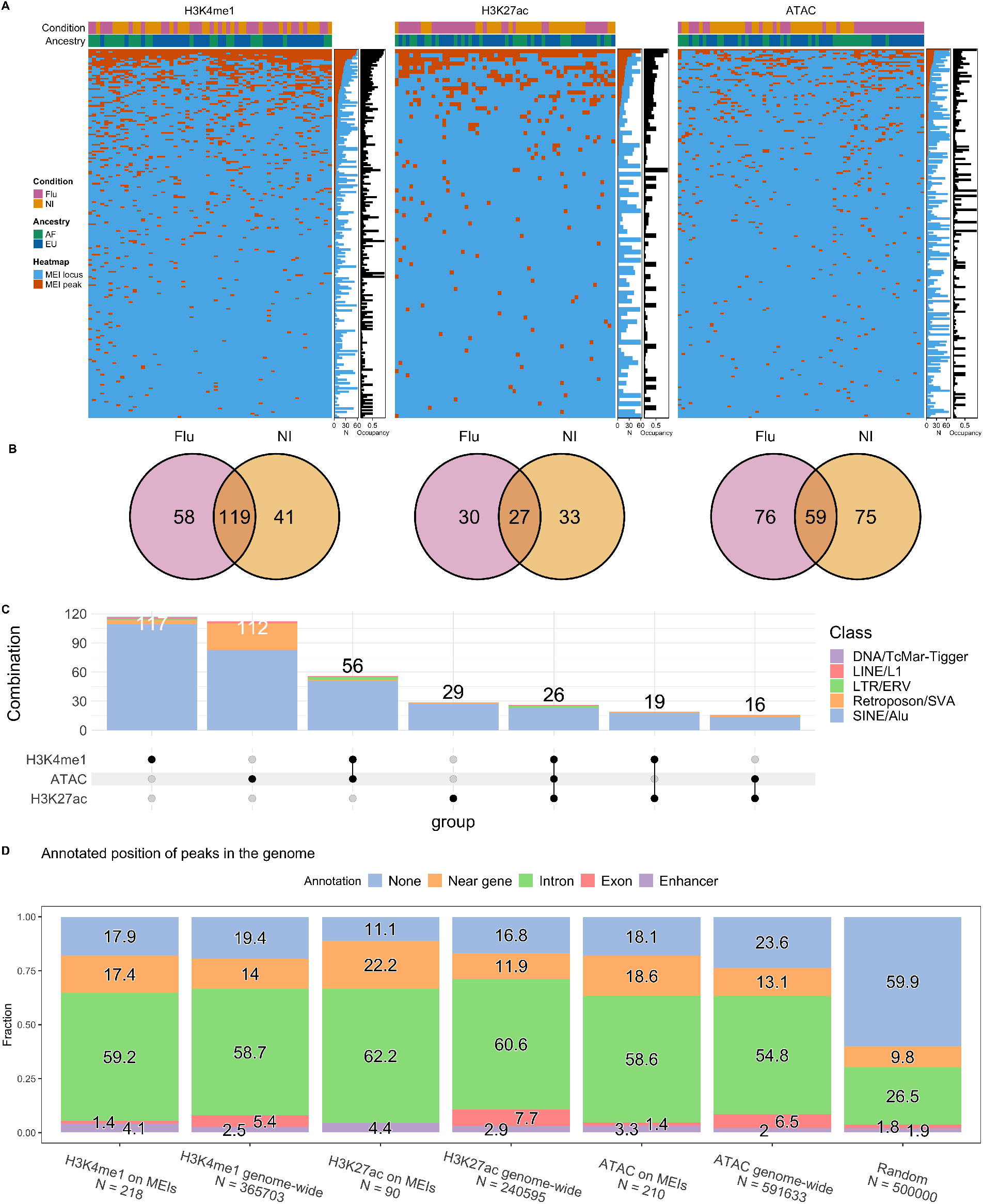
Cohort data reveals MEIs that act as potential enhancers. A) Summary of MEIs (rows) that support H3K4me1, H3K27ac and ATAC peaks in the cohort samples (columns). Occupancy is the ratio between samples that support peaks on the MEI (N - red) and those that carry the MEI (N - blue). B) Venn diagrams showing MEI peaks that are shared between flu-infected and non-infected conditions. C) Upset plot describing the number of MEIs that support a combination of H3K4me1, H3K27ac and ATAC peaks, annotated by transposable element class. D) The annotated positions of genome-wide peaks and MEI peaks in the genome, with uniformly and randomly sampled genome positions for comparison. Peaks near genes are within 10 Kbp of a gene boundary.

Next, we looked for MEIs that carry combinations of epigenomic marks that are characteristic of enhancer sequences (Fig 5C). For example, 56 MEIs supported both H3K4me1 and ATAC, a combination which suggests poised enhancers in open chromatin. Similarly, 16 instances showed H3K27ac and ATAC and 26 insertions were marked by H3K4me1, H3K27ac and ATAC, which is strong evidence for active enhancers in open chromatin. There is a noticeable number of SVA elements showing ATAC peaks, second only to Alu elements. We find that peaks on MEIs have a similar distribution relative to exons, introns, genes and enhancers as compared to other peaks and quite distinct from a random distribution (Fig 5D). Additionally, we find dozens of MEI loci that are in the vicinity of genes associated with “viral processes”, “immune system processes”, the “inflammatory response” and the “positive regulation of NF-kappaB” (Fig S14). For example, an Alu insertion that supports H3K27ac peaks (Fig S15A) in 21 samples is immediately upstream of *CD300E*, an immune-activating receptor gene [35]. This frequent MEI peak is also located within DNase and transcription factor clusters (Fig S15B), which is further evidence for active chromatin.

Finally, since Alu polymorphisms could alter gene transcript levels [36], we asked if any MEIs were eQTLs for expressed genes. We mapped MEI-eQTLs and found 18 MEIs in the flu-infected condition and 34 MEIs in the non-infected condition that were associated with gene expression (FDR lower than 5 × 10^−2^, Methods). In total, there are also 354 MEIs that are ca/hQTLs for at least one mark or condition (Fig S16). Of the MEI-eQTLs, 3 have a supporting caQTL or hQTL in the flu-infected condition and 9 in the non-infected condition (Table S2). In particular, in the flu-infected condition, we detected an AluYh3 MEI that is an eQTL for *DGKE* and *TRIM25*, a gene that restricts influenza RNA synthesis [37], and is also a QTL for 2 H3K4me1 peaks, 2 H3K27ac peaks and 3 ATAC peaks (Table S3). We show the average read depth at this locus in flu-infected samples that carry the MEI (Fig 6A), the flu-infected samples that do not carry the MEI (Fig 6B), in non-infected samples that carry the MEI (6C) and in non-infected samples that do not carry the MEI (Fig 6D). The more traditional linear projection of the average read depth at this locus confirms that the signal is higher in flu-infected samples that carry the MEI when using an accurate genome graph (Fig 6E-F). Overall, this MEI exists in a flu-specific active chromatin state, is associated with *TRIM25* and *DGKE* gene expression and would have been missed with the linear reference genome.

**Figure 6:**
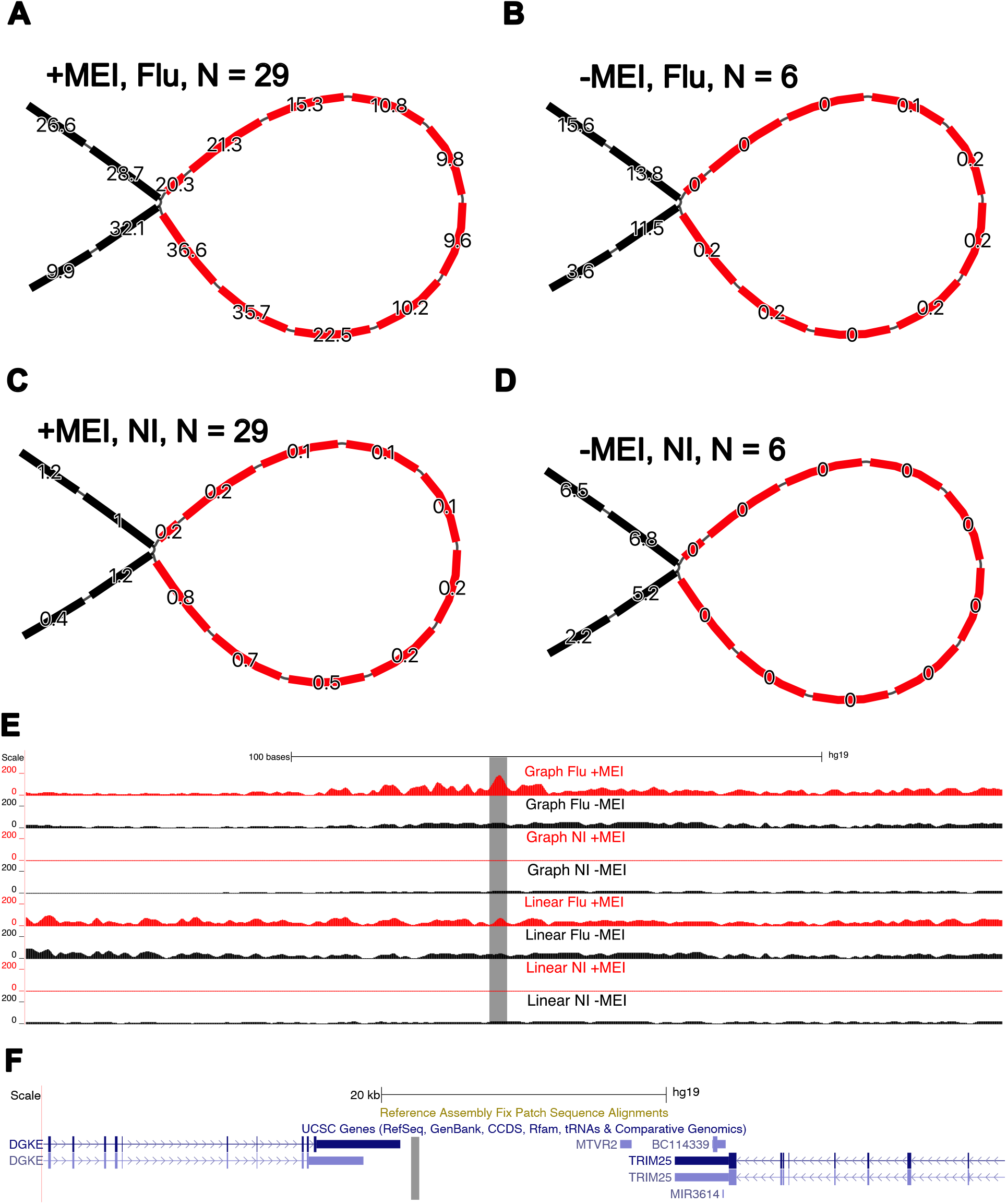
Example of flu-specific and genotype-specific peak on an MEI. A) Average H3K27ac read depth in the locus of an AluYh3 MEI-eQTL in flu-infected samples that carry the insertion, B) in flu-infected samples that do not carry the MEI, C) in non-infected samples that carry the MEI and D) in non-infected samples that do not carry the MEI. The read depths of homozygous nodes were halved before averaging. Reads below a MAPQ of 10 were not counted. E) A genome browser view of the read depth after projecting alignments onto the linear genome, contrasting the alignments to the graph genome and the reference genome. F) Nearby genes that are associated with this MEI-eQTL (DGKE, TRIM25). The grey strips denote the position of the MEI.

## Discussion

We have constructed a genome graph to encapsulate the genetic diversity of a cohort of 35 individuals by re-sequencing their genome with linked-reads. These data allowed us to reliably discover and phase SNVs, small indels and a large number of MEIs; the latter through the development of a local assembly method. We note that, because of the limitations of the linked-read technology, the MEIs included were skewed towards the shorter insertions and missed a significant portion of larger events. Indeed, out of the 1344 L1 polymorphisms detected in the cohort, 1048 (78%) could not be assembled and integrated into the genome graph. We anticipate that long read technologies [25] will be needed to construct more complete genome graphs that represent even more complex SVs.

We showed that our cohort genome graph improves read mapping for epigenomic data. This lead to a net increase in the number of peak calls, which are replicated across samples and occur in functionally interesting loci such as in the vicinity of immune genes. In addition, we showed that using a genome graph had an impact on the discovery of histone and chromatin accessibility QTLs in polymorphic regions. However, these new results were restricted to regions where sequence variants were added the graph, which based on our current approach remain relatively small (SNPs, indels and short MEIs). Therefore, the gains were mostly associated smaller epigenomic features such as narrow histone marks or chromatin accessibility. We anticipate that adding larger and more complex structural variations to the graph, such as segmental duplications, will reveal even more epigenomic features that are present but difficult to measure using traditional approaches.

Furthermore, we used the cohort genome graph to re-genotype 5598 MEIs using short read WGS data and reliably assign epigenomic signal to MEI alleles. While rare, we sometimes observed heterozygous MEIs where the majority of the epigenomic signal was either on the insertion or the reference allele, revealing homologous loci that are in different chromatin states. This shows that genome graphs can be exploited to search for epigenomic signal that is specific to alleles in a cohort of individuals. Meanwhile, current methods that measure allelic specific signal are limited in their ability to represent complex structural variants or relate genomes within a population to one another. Therefore, genome graphs could prove a powerful framework to study the chromatin state of polymorphic structural variants in a large number of genomes.

Finally, with the recent completion of the first full human genome [38], there is a general appreciation of the need to use a new baseline for genomic and epigenomic analysis, including graph genomes that can capture genetic diversity. The Human Pangenome Reference Consortium is making great strides towards this goal [39] and new approaches are needed to demonstrate the benefits of these richer references. Once a human pan-genome graph is developed from a set of high quality and diverse genome assemblies, a more comprehensive set of structural variants will become easily accessible. The hope is that similar analyses as was done here could be employed on a multitude of cell types and phenotypes to identify functionally relevant SVs without the challenges associated with the construction of cohort-specific genome graphs.

## Methods

### ATAC-seq and ChIPmentation library preparation

ATAC-Seq library preparation was performed according to the Omni-ATAC protocol [40]. 50,000 GM12878 cultured cells were resuspended in 1 ml of cold ATAC-seq resuspension buffer (RSB; 10 mM Tris-HCl pH 7.4, 10 mM NaCl, and 3 mM MgCl2 in water). Cells were centrifuged at 500 g for 5 min in a pre-chilled (4 °C) fixed-angle centrifuge. After centrifugation, supernatant was aspirated and cell pellets were then resuspended in 50 *µ*l of ATAC-seq RSB containing 0.1% IGEPAL, 0.1% Tween-20, and 0.01% digitonin by pipetting up and down three times. This cell lysis reaction was incubated on ice for 3 min. After lysis, 1 ml of ATAC-seq RSB containing 0.1% Tween-20 (without IGEPAL and digitonin) was added, and the tubes were inverted to mix. Nuclei were then centrifuged for 10 min at 500 rcf in a pre-chilled (4 °C) fixed-angle centrifuge. Supernatant was removed and nuclei were resuspended in 50 *µ*L transposition mix (2x TD Buffer, 100 nM final transposase, 16.5 *µ*L PBS, 0.5 *µ*L 1% digitonin, 0.5 *µ*L 10% Tween-20, 5 *µ*L H2O). Transposition reactions were incubated at 37 °C for 30 min in a thermomixer with shaking at 1000 rpm. Reactions were cleaned up with Zymo DNA Clean and Concentrator 5 columns. Primers (i5 and i7) were added by amplification (12 cycles) using NEBNext 2x MasterMix. Sequencing of the ATAC-Seq libraries was performed on the Illumina NovaSeq 6000 system using 100-bp paired-end sequencing.

ChIPmentation library preparation was performed on 5 million cross-linked cells (1% formaldehyde). After cell lysis, sonication of nuclei was performed on a BioRuptor UCD-300 targeting 150-500 bp size. Immunoprecipitation and library preparation for the histone marks H3K27ac and H3K4me1 was performed following the Auto-ChIPmentation protocol for Histones (Diagenode inc, Denville, USA) according to the manufacturer’s indications. Sequencing of the ChIPmentation libraries was performed on the Illumina NovaSeq 6000 system using 100-bp paired-end sequencing.

### Whole genome sequencing with linked reads and genotyping

High molecular weight DNA was extracted using the MagAttract HMW kit from Qiagen and quantified using the Qubit dsDNA HS assay kit (ThermoFisher) for 35 samples. In order to generate the linked read libraries 1ng of HMW DNA was loaded on a 10x Chromium device using 10x Genome Sequencing Solution v2 reagents (10X Genomics). Illumina compatible libraries were prepared as per the 10x Genomics protocol and loaded as one library per lane on the Illumina HiSeqX instrument for 150bp paired-end sequencing. SNVs and indels were called using the `longranger` pipeline.

### Locally assembling linked reads with BarcodeAsm

For the purpose of locally assembling linked reads, we wrote BarcodeAsm [41]. The inputs to BarcodeAsm are a BED file describing the regions to be locally assembled and a BAM file that was aligned with `lariat` [17]. This BAM file must be provided twice, once sorted by position and once sorted by barcode (with `bxtools`).

BarcodeAsm uses the position sorted BAM file to identify barcodes that are present in the target assembly window. Then it moves to the barcode sorted BAM file to retrieve all the reads that are tagged by the previously identified barcodes.

BarcodeAsm also allows filtering reads recovered from outside the local window (800 bp for Alus, 8000 bp for other TEs) by mapping quality. We introduced this feature because we expect reads that belong to novel transposable element insertions to be unmapped or mapped to the wrong copy, have very low mapping quality, or be multi-mapped.

Next, the `fermi-lite` [42] library assembles the resulting collection of reads and creates a unitig graph, from which the contigs associated with the assembly window are extracted. Finally, BarcodeAsm aligns the contigs to the local window with `minimap2` [43]. A select set of fermi-lite assembly parameters and `minimap2` alignment parameters are exposed via the command line and can be adjusted to each application.

### Reassembling transposable element insertions

ERVcaller [44] and MELT [45] were used to genotype novel insertions of Alu, LINE1, SVA, and ERV transposable elements using short reads. To further recover potential insertions within the same type of reference repeats (nested TE insertions), candidate nested insertions that were detected in other public datasets were not removed. For each genotype, local genomic windows were centered on the insertion site. These windows are 800 bp in length for the short Alu elements and 8 Kbp for the longer transposable elements. To optimize the outcomes of the assembly, BarcodeAsm was run separately on the short (≈ 300 bp) and long insertions (> 300 bp) with different parameters. For Alus, a minimum read overlap of 30 bp, a maximum mapping quality of 10 (as assigned by `lariat`), and a minimum k-mer frequency of 2 were required. For the larger MEIs, a minimum read overlap of 35, a maximum mapping quality of 20, and a minimum k-mer frequency of 8 were used instead. These parameters were found through a grid search approach and are expected to vary with different data sets.

For the NA12878 benchmark, the SRA accession numbers ERR174324, ERR174325 to ERR174341 were merged in order to genotype MEIs. MEIs were assembled from the public NA12878 linked reads hosted at https://support.10xgenomics.com/genome-exome/datasets/2.0.0/NA12878_WGS. Insertions were extracted directly from the BarcodeAsm output using `scripts/alignment to vcf.py` to create a VCF file.

For the larger cohort, a consensus sequence approach was used to take advantage of multiple MEI copies in the population (see `BarcodeAsm/scripts/`). Here, contigs that contain insertions are selected but not immediately used to recover an insertion. Instead, `scripts/msa.py` generates a multiple alignment using MUSCLE and calculates a consensus contig for a particular locus across samples. This consensus contig is aligned back to the local window to call the consensus insertion and to create a multi-sample VCF file using `scripts/extract consensus.py`. For NA12878, the assembled contigs were validated by matching them against its haplotype resolved de novo assembly [24] using `minimap2 -H` [43] and selecting the haplotype with the best mapping quality.

### Annotating assembled insertions

To annotate the insertions (without any flanking sequences), we used RepeatMasker with the Dfam [46] database and the longest annotation was selected for each insertion. For each assembled TE polymorphism, the distance to the nearest enhancer in the GeneHancer annotation [47] and the distance to the nearest exon in the GENCODE annotation [48] were calculated. The same was done with the MEIs from the 1000 Genome Project [32] and with an equal number of random positions sampled uniformly from the genome. The population structure of the MEI genotypes were compared to the population structure of WGS variants by running principal component analysis with `SNPRelate` [49].

### Creating and benchmarking genome graphs

The genomes graphs were generated with `vg construct` [26] on VCF file containing SNVs, indels and MEIs. For the benchmark genome graph, the NA12878 Platinum callset [50] was merged with the MEI VCF file. For the cohort genome graph, SNPs and indels were called using the LongRanger pipeline [21] independently for each sample were merged with the population MEIs to create a multi-sample VCF file. A Nextflow script to generate the genome graphs from these inputs and the b37 reference genome is found in `pop graph.nf`. The sensitivity and specificity of the resulting genome graphs was checked by aligning a matching but separate WGS data set of the same samples (downsampling the merged read set by 5x in the case of NA12878) and removing non-specific alignments with `vg filter -r 0.90 -fu -m 1 -q 10 -D 999`. After, the assembled MEI snarls were genotyped with `vg call -m2,4` [51] and the graph genotypes were compared to ERV and MELT genotypes.

To evaluate the impact of the genome graphs on alignment, a WGS read set was simulated from the diploid assembly of NA12878. To achieve this, a genome graph was created from hg19 using the structural variant sequences called by the authors directly from the chromosome scale assembly. Paired-end reads were simulated using `vg sim` from this graph with a fragment size of 2000 bp, to ensure that we can access at least some of the long copy number variants that have low mappability. First, this read set was aligned to the hg19 reference graph. Then, the alignment of the simulated reads was repeated on increasingly complete genome graphs, first by including only SNVs, then indels and lastly MEIs. Finally, the simulated reads were aligned to the NA12878 de novo genome graph, which is the true genome by construction. The previous alignments are compared against this last alignment using `vg gamcompare`.

### Evaluating the impact of genome graphs on peaks

In parallel, ChIP-seq and ATAC-seq data was aligned to the reference graph to obtain reference peaks. The reference peaks were intersected with the graph peaks, and categorized into common peaks, graph-only and ref-only peaks. Common peaks are unaltered peaks that are found with both the reference and cohort genome graph. Graph-only peaks and ref-only peaks are altered peaks that are only found in the cohort graph or the reference graph respectively. A logistic regression model including peak width, the presence of an indel and the presence of a SNP was fitted using `cv.glnmnet` (see `peak variants.R`) [52]. The number of common and altered peaks were balanced by subsampling common peaks. We report the cross-validation mean coefficients and median AUC. To summarize the population properties of peaks, curves were generated for common, graph-only and ref-only peaks (see `Rscripts/peak replication.R`). The width of each peak was fixed to 200 bp and the proportion of samples that have a peak at the same location was calculated. The resulting curve is the inverse cumulative distribution of peak frequencies. A permutation simulation was performed to obtain the inverse cumulative distribution that is expected from random overlaps. In the simulation, a new peak set of the same size is randomly sampled for each individual from the set of all peaks and the overlaps are used to recompute the inverse cumulative distribution. The simulation was run 100 times and the average inverse cumulative distribution is reported.

### QTL mapping

We filtered to exclude non-autosomal and non-biallelic variants. Additionally, we removed SNPs and MEIs that had a call rate of *<*90% across all samples, that deviated from Hardy–Weinberg equilibrium at *p <* 10^−5^, and with minor allele frequency less than 5%. This resulted in 7,383,243 SNPs and 1222 MEIs used for QTL mapping. We used the R package `MatrixeQTL` [53] to examine the associations between SNP genotypes and chromatin accessibility, H3K4me1 and H3K27ac histone marks using both graph and reference read counts. We also mapped MEI-eQTLs to find association between mobile element insertion genotypes and gene expression counts (derived from alignments to the reference genome with `STAR` [54]). In each case, we calculated age and batch corrected expression matrices. Here, batch is a categorical variable. We calculated normalization factors to scale the raw library sizes using `calcNormFactors` in `edgeR (v 3.28.1)` [55] and used the `voom` function in `limma (v 3.42.2)` [56] to apply these factors to estimate the mean-variance relationship and convert raw read counts to *logCPM* values. We then fit a model using mean-centered age and admixture, removing batch effects using `ComBat` from the `sva` Bioconductor package [57]. We then regressed out age effects, resulting in the age and batch corrected expression matrices used as inputs for `MatrixeQTL`. To increase the power to detect cis-QTL, we accounted for unmeasured-surrogate confounders by performing principal component analysis (PCA) on the age and batch corrected expression matrices. The number of PCs chosen for each data type empirically led to the identification of the largest QTL in each condition and are reported in Table S4.

Mapping was performed combining individuals in order to increase power, thus, we included the first eigenvector obtained from a PCA on the SNP data as a covariate in our linear model in order to account for population structure. Local associations (i.e., putative cis QTL) were tested against all SNPs and MEIs located within the peak or gene or 100 Kbp upstream and downstream of each peak or gene. We recorded the strongest association (minimum p-value) for each peak/gene, which we used as statistical evidence for the presence of at least one QTL for that peak/gene. We permuted the genotypes ten times, re-performed the linear regressions, and recorded the minimum p-value for each peak/gene for each permutation. We used the R package `qvalue` [58] to estimate FDR. In all cases, we assume that alleles affect phenotype in an additive manner.

### Functional enrichment analysis

To find if peaks are enriched in functional pathways, the names of genes that are within 10 Kbp of a peak were compiled into gene sets (see `pathways.R`). A gene may appear only once in a gene set, even if it is nearby several peaks. This was done separately for the flu-infected and non-infected conditions and for common and graph-only H3K4me1, H3K27ac and ATAC peaks. Peaks that occur only in one sample within a condition were excluded. In a separate but similar analysis, we compiled gene sets that are within 10 Kbp of a peak that is under the control of a QTL, separately for H3K4me1-QTLs, H3K27ac-QTLs and ca-QTLs, and separately in the infected and non-infected condition.

Then, gene sets were checked for functional enrichment [59, 60] in Biological Process Gene Ontology (GO:BP) terms with `gprofiler2` [61]. We also calculated the gene ratio of enriched terms, which is the share of genes in the list associated with a GO term.

### Evaluating peaks on genome graphs

H3K27ac, H3K4me1 ChIP-seq and ATAC-seq libraries were aligned to the genome graphs and peaks were called with Graph Peak Caller [27] to obtain graph peaks. All the snarls in the genome graph were genotyped with `vg call -m2,4` in order to partition the reads between the reference and alternative alleles in peaks that overlap a polymorphism. This information is available in the AD (alternate allele depth) and DP (site depth) tags in the VCF output. As before, non-specific alignments were removed. This approach was validated by listing peaks that overlap polymorphic loci in homozygous reference samples to create a negative control peak set, which should not show any coverage on the alternate allele.

Since the MEI allele is longer than the reference allele, we scaled down read counts (see `Rscripts/allele support.R`) on the longer allele according to:.

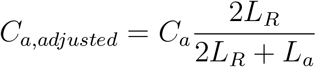

where *C*_*a*_ denotes the read count on the long allele, *L*_*R*_ the read length and *L*_*a*_ the length of the long allele.

Further, peaks were binned according to the partitioning of reads between the reference and alternative alleles using a two sided binomial test with *α* = 0.05. Peaks that were skewed towards the MEI allele (p-value ≤ *α/*2) are placed in the MEI support bin. Peaks that show roughly equal read coverage on both alleles (p-value ≥ *α/*2) are labeled as biallelic. When reads are skewed towards the reference allele (p-value ≤ *α/*2), the peak belongs to the reference support bin. We also counted the MEIs that support combinations of marks to support their functional relevance [62, 63, 64, 65].

## Ethics approval and consent to participate

Not applicable.

## Consent for publication

Not applicable.

## Availability of data and materials

- The source code of BarcodeAsm is available on the Zenodo repository [41].
- Additional code and processed data to reproduce the analysis, figures and manuscript are available on the Zenodo repository [66].
- Code for QTL mapping is available on the Zenodo repository [67].

## Additional files

## Supplementary Tables

**Table S1:** Relative influence of peak width (100bp), SNPs and indels on the log-odds of a peak being found only with the cohort graph genome.

| Peak | snp | indel | width | intercept | AUC |
| --- | --- | --- | --- | --- | --- |
| H3K4me1 | 0.11 | 0.55 | -0.72 | 2.44 | 0.91 |
| H3K27ac | 0.12 | 0.56 | -0.78 | 2.72 | 0.92 |
| ATAC | 0.19 | 0.45 | -0.87 | 2.70 | 0.90 |

**Table S2:** MEI-eQTLs that support epigenomic marks and are associated with gene expression at a false discovery rate below 5 × 10^−2^ in the flu-infected or the non-infected condition.

| MEI-eQTL | Condition | Marks | Gene | Beta | FDR |
| --- | --- | --- | --- | --- | --- |
| chr1:160047665 | NI | H3K4me1, ATAC | ENSG00000177807 | 1.02 | $4.10 \times 10^{-2}$ |
| chr5:1812640 | NI | H3K4me1 | ENSG00000171421 | 0.17 | $2.48 \times 10^{-3}$ |
| chr5:78442870 | Flu | H3K4me1 | ENSG00000152409 | 0.26 | $4.01 \times 10^{-2}$ |
| chr7:128209146 | NI | H3K4me1, ATAC | ENSG00000242588 | 0.15 | $1.13 \times 10^{-3}$ |
| chr7:128217333 | NI | ATAC | ENSG00000242588 | 0.16 | $3.87 \times 10^{-4}$ |
| chr8:42039896 | NI | H3K4me1, H3K27ac, ATAC | ENSG00000070718 | 0.15 | $4.12 \times 10^{-3}$ |
| chr11:47806640 | NI | H3K27ac | ENSG00000030066 | 0.28 | $3.99 \times 10^{-7}$ |
| chr11:47806640 | NI | H3K27ac | ENSG00000252874 | 0.41 | $1.06 \times 10^{-2}$ |
| chr12:124066475 | NI | H3K4me1, ATAC | ENSG00000111364 | -0.19 | $1.13 \times 10^{-3}$ |
| chr14:92586933 | NI | H3K4me1 | ENSG00000183648 | -0.08 | $4.44 \times 10^{-2}$ |
| chr14:92586933 | Flu | H3K4me1 | ENSG00000165934 | -0.15 | $9.45 \times 10^{-4}$ |
| chr15:41100318 | NI | H3K4me1, H3K27ac, ATAC | ENSG00000166140 | -0.11 | $9.58 \times 10^{-3}$ |
| chr17:54947569 | Flu | H3K4me1, H3K27ac, ATAC | ENSG00000153933 | 0.91 | $9.56 \times 10^{-16}$ |
| chr17:54947569 | Flu | H3K4me1, H3K27ac, ATAC | ENSG00000214226 | 0.44 | $9.69 \times 10^{-9}$ |
| chr17:54947569 | Flu | H3K4me1, H3K27ac, ATAC | ENSG00000121060 | 0.24 | $2.13 \times 10^{-8}$ |
| chr20:13704145 | NI | H3K4me1, ATAC | ENSG00000101247 | 0.14 | $3.69 \times 10^{-4}$ |

**Table S3:** Summary of MEI-QTLs listing the regressed features (peaks within 10 Kbp or gene), the biological condition, the effect size, and the false discovery rate. Each entry describes the insertion coordinate, the marks supported by the MEI and the TE family.

| QTL | Effect size | FDR |
| --- | --- | --- |
| chr17:54947569 — H3K4me1, H3K27ac, ATAC — AluYh3 |  |  |
| *TRIM25* - Flu | 0.239 | $2.13 \times 10^{-8}$ |
| *DGKE* - Flu | 0.912 | $9.56 \times 10^{-16}$ |
| H3K4me1:chr17:54944305-54949599 - Flu | 0.435 | $1.68 \times 10^{-5}$ |
| H3K4me1:chr17:54952157-54958631 - Flu | 0.114 | $2.79 \times 10^{-2}$ |
| H3K27ac:chr17:54945079-54949298 - Flu | 0.507 | $8.85 \times 10^{-7}$ |
| H3K27ac:chr17:54956812-54958346 - Flu | 0.646 | $6.95 \times 10^{-5}$ |
| ATAC:chr17:54946838-54948880 - Flu | 0.923 | $3.85 \times 10^{-12}$ |
| ATAC:chr17:54945058-54946521 - Flu | 0.466 | $1.67 \times 10^{-8}$ |
| ATAC:chr17:54952726-54958278 - Flu | 0.252 | $1.15 \times 10^{-5}$ |
| chr11:47806640 — H3K27ac — AluYh7 |  |  |
| *NUP160* - NI | 0.278 | $3.99 \times 10^{-7}$ |

**Table S4:**
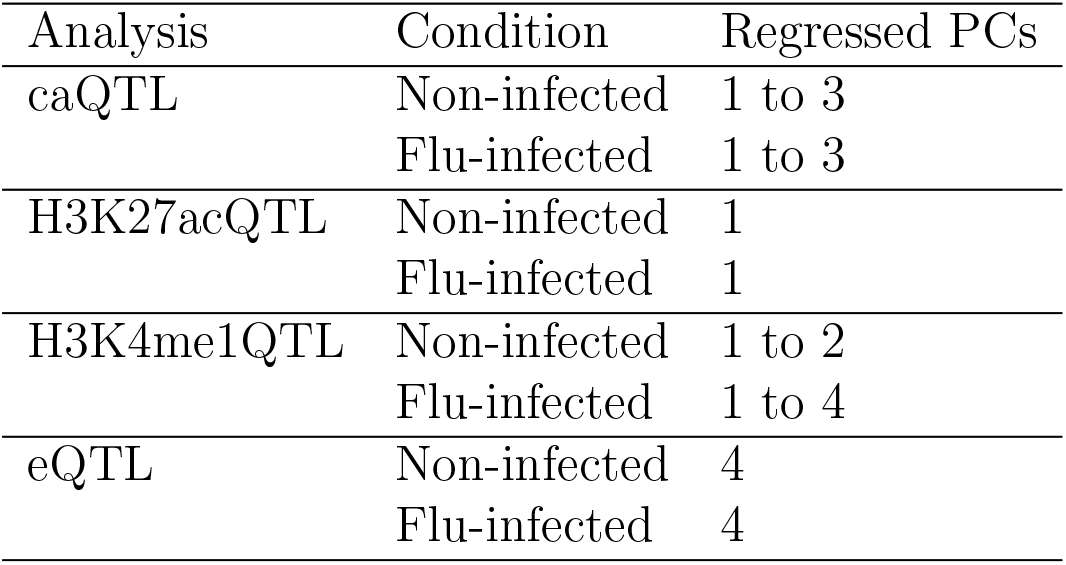
Number of principal components performed on age and gene expression included in each QTL discovery analysis in order to correct for age and batch effects.

## Supplementary Figures

**Figure S1:**
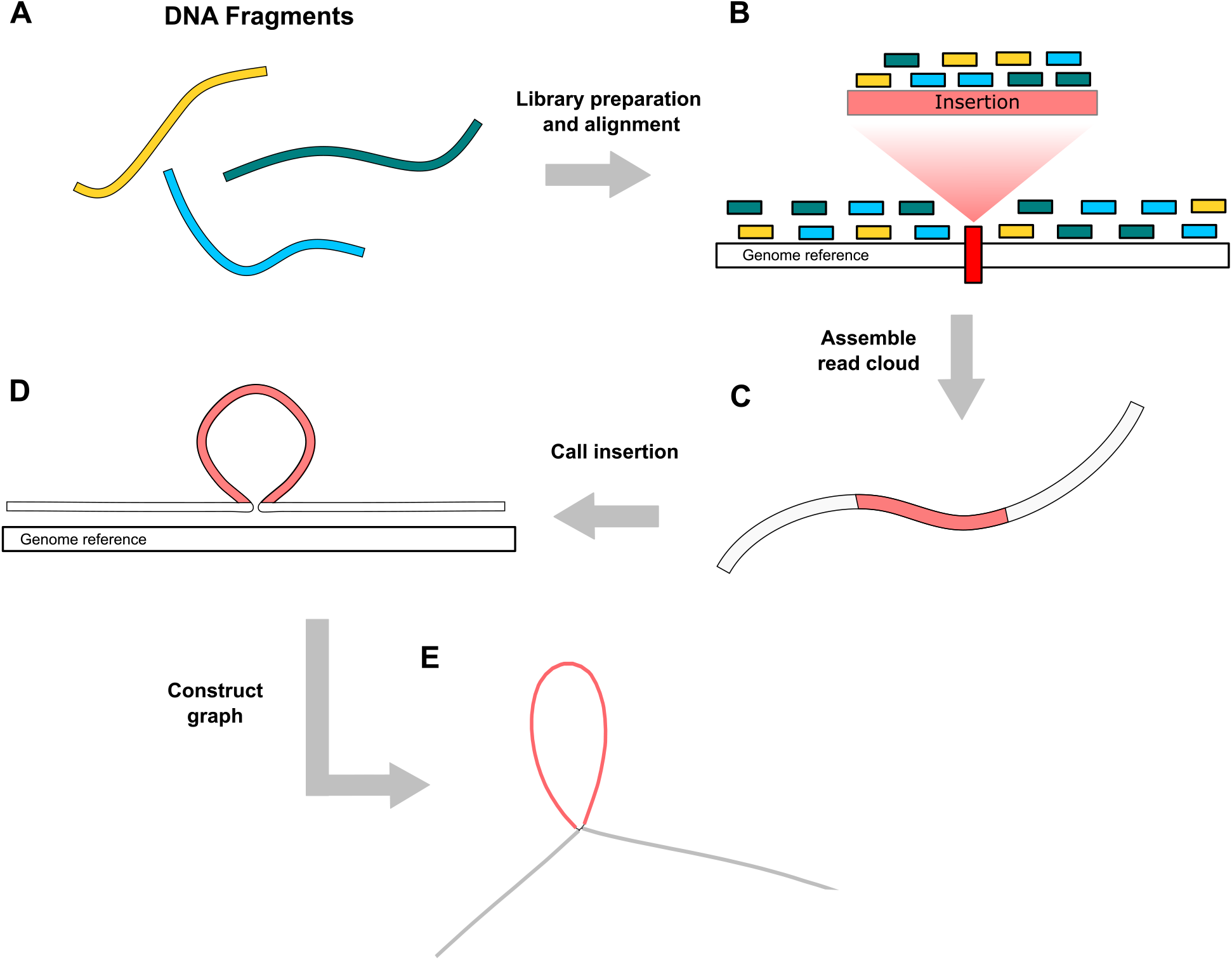
A) DNA fragments that are associated with barcodes (yellow, blue, green). B) Throughout the sequencing library protocol and alignment to the reference genome with lariat, the short reads remain associated with the barcode of the source fragment. We enumerate the barcodes observed around an insertion site. C) We assemble the reads tagged by the previously enumerated barcodes site with fermi-lite to obtain a contig that covers the insertion (red). D) We realign the contig back to the genomic locus to identify the boundaries of the inserted sequence. E) We augment the genome graph with a bubble that represents the assembled insertion.

**Figure S2:**
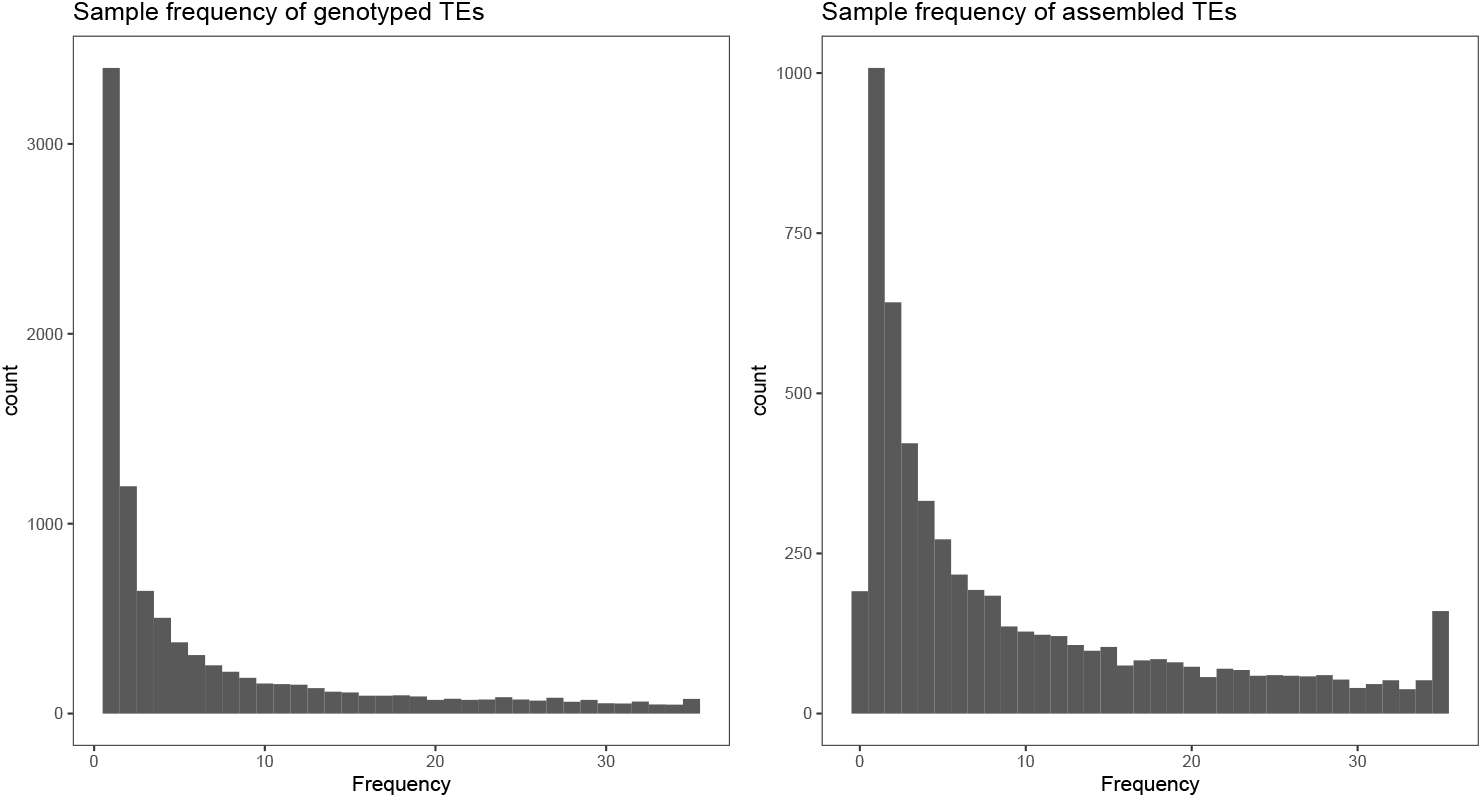
Population frequency of the A) genotyped and B) assembled MEIs in the cohort.

**Figure S3:**
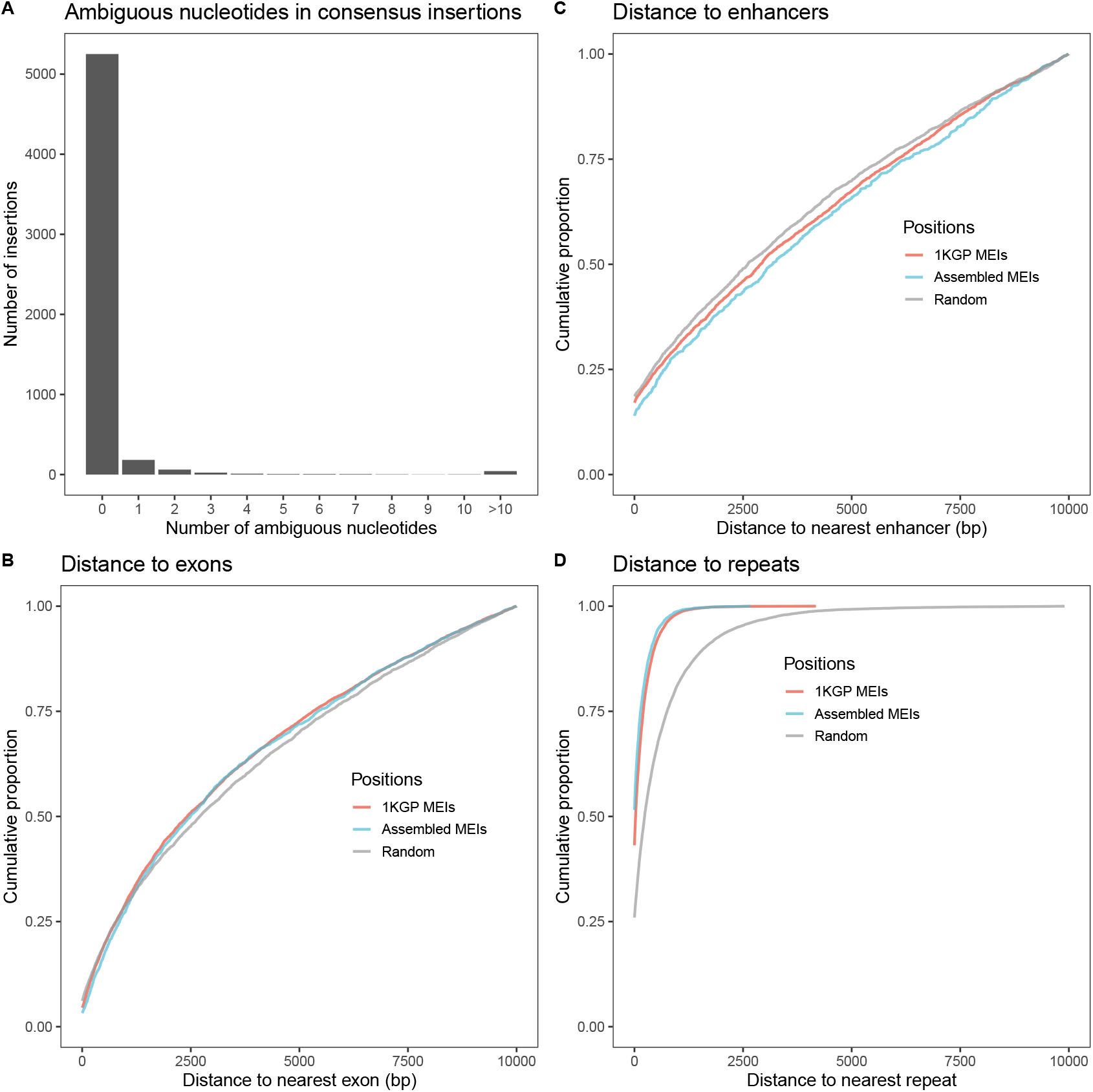
A) Number of ambiguous nucleotides per sequence in consensus insertions. The distributions of insertions in the genome around B) exons, C) enhancers and D) other repeats.

**Figure S4:**
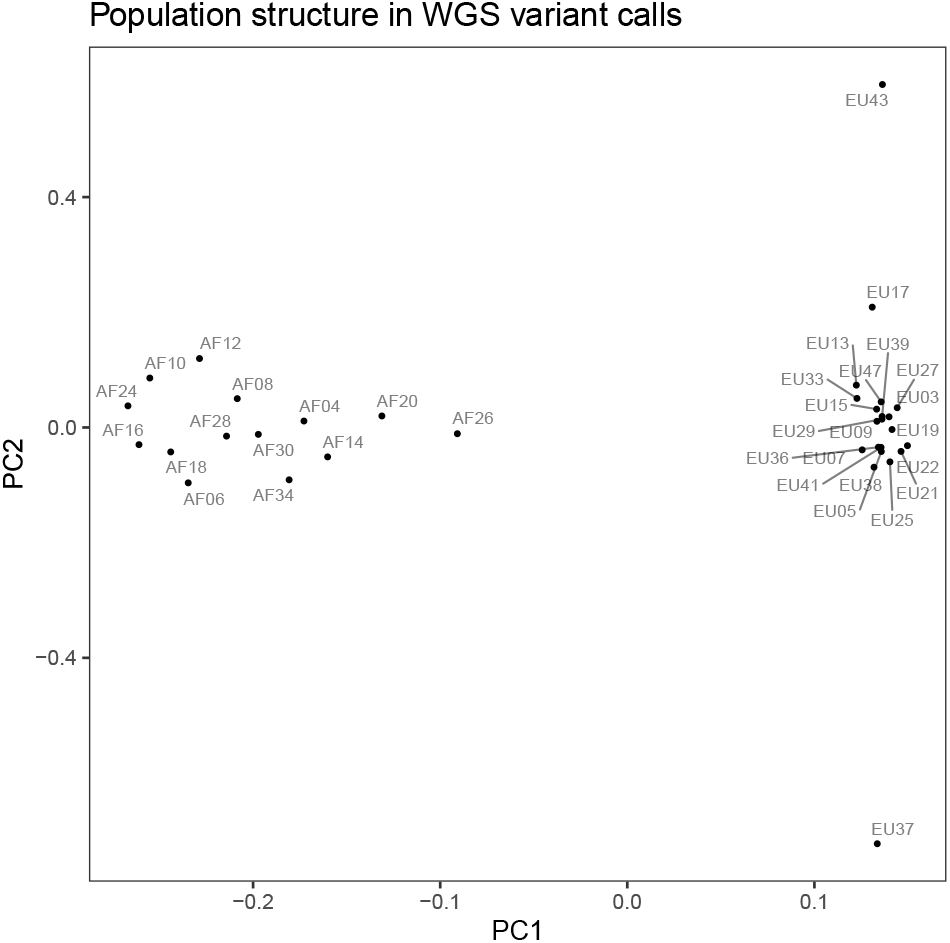
PCA projection showing the population structure observed in whole genome sequencing genotypes.

**Figure S5:**
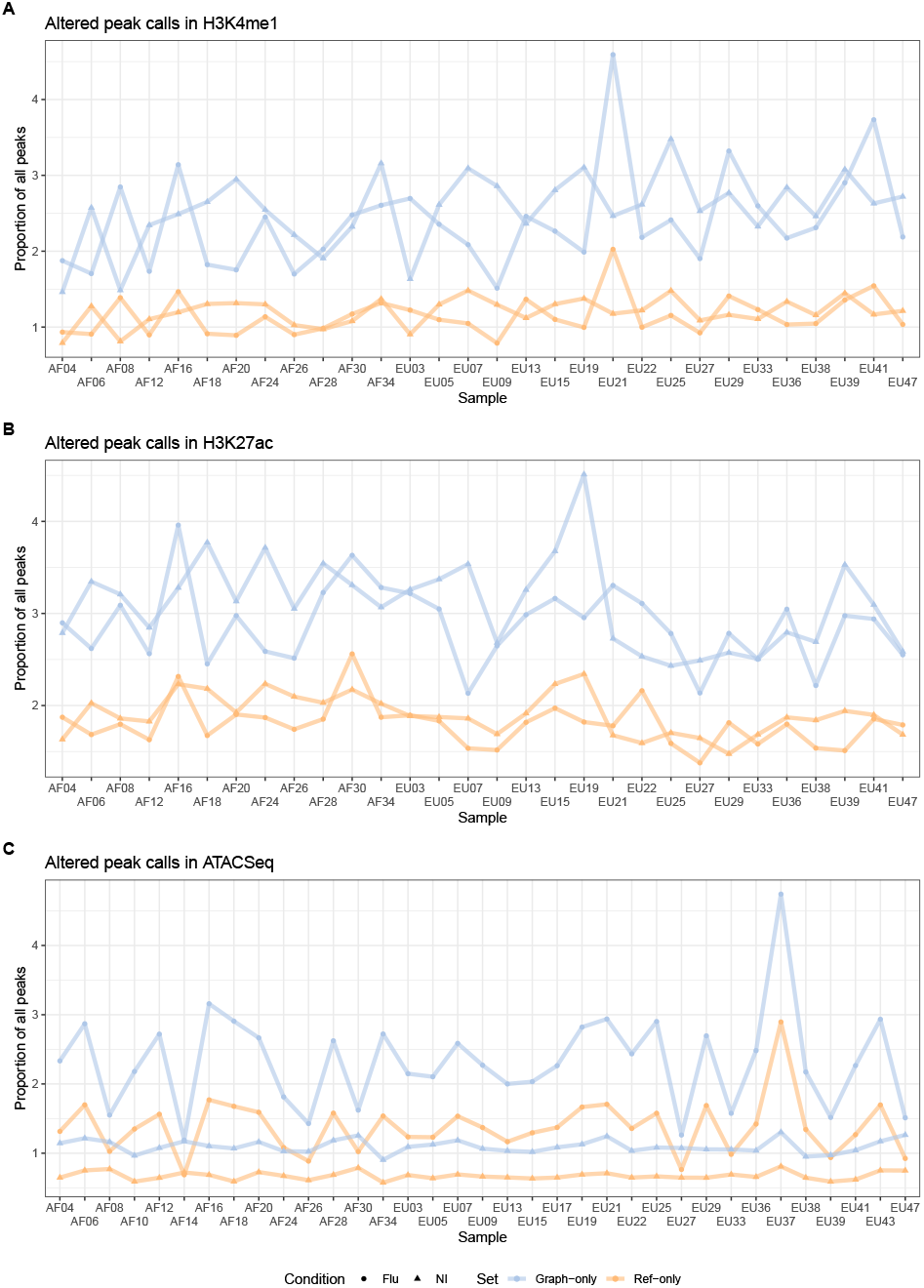
Frequency of altered peak calls as percentages of all peaks in the sample.

**Figure S6:**
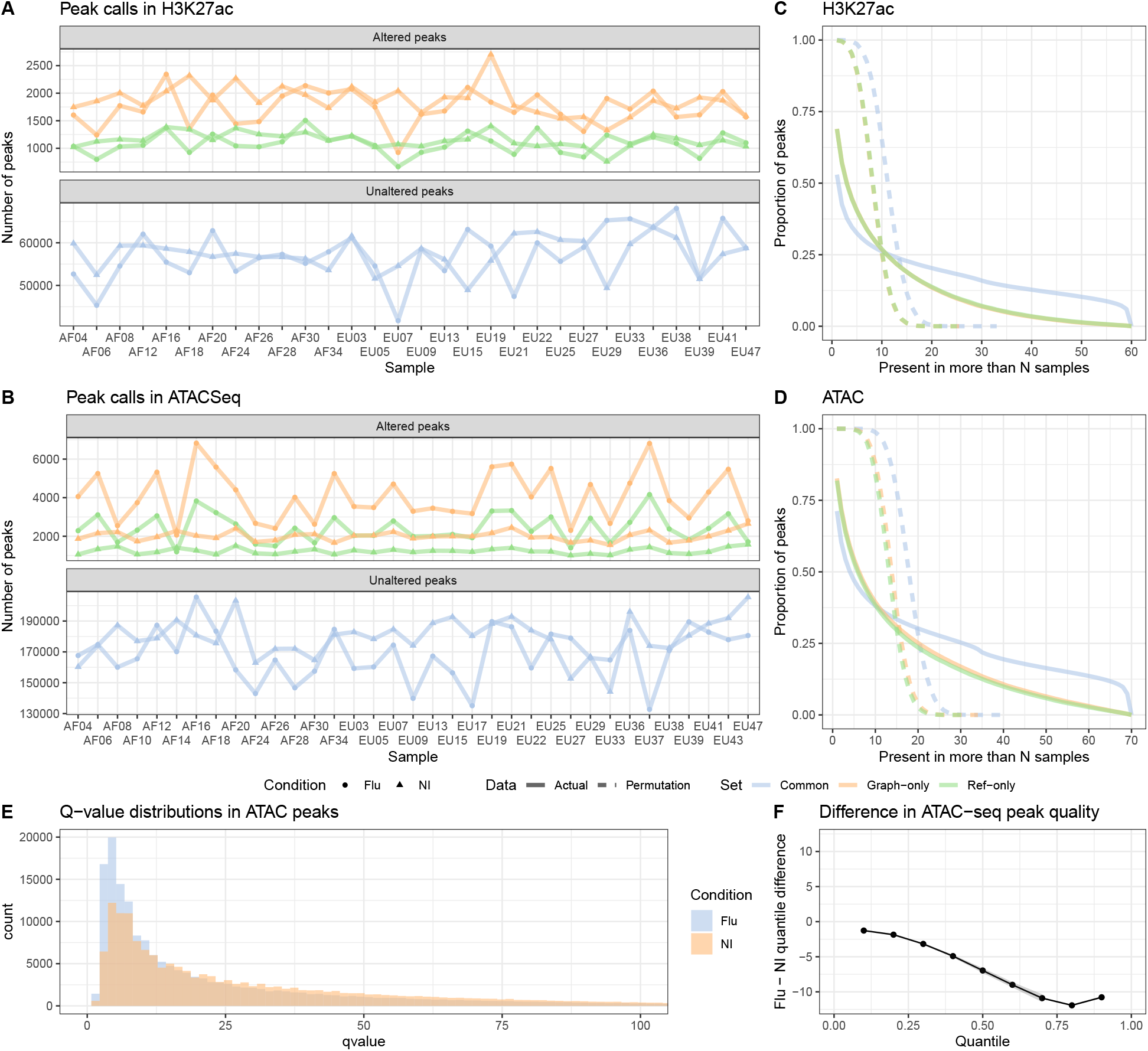
A) B) The number of altered (graph-only, ref-only) and unaltered (common) peaks between the cohort and the reference genome graphs for H3K27ac and ATAC-seq. C) D) Inverse cumulative distributions describing how many H3K27ac and ATAC-seq peaks are observed in more than a number of samples E) Q-value distribution of ATAC peaks in flu-infected and non-infected samples. F) The shift function of the two distributions, describing the difference in q-values at the same quantiles.

**Figure S7:**
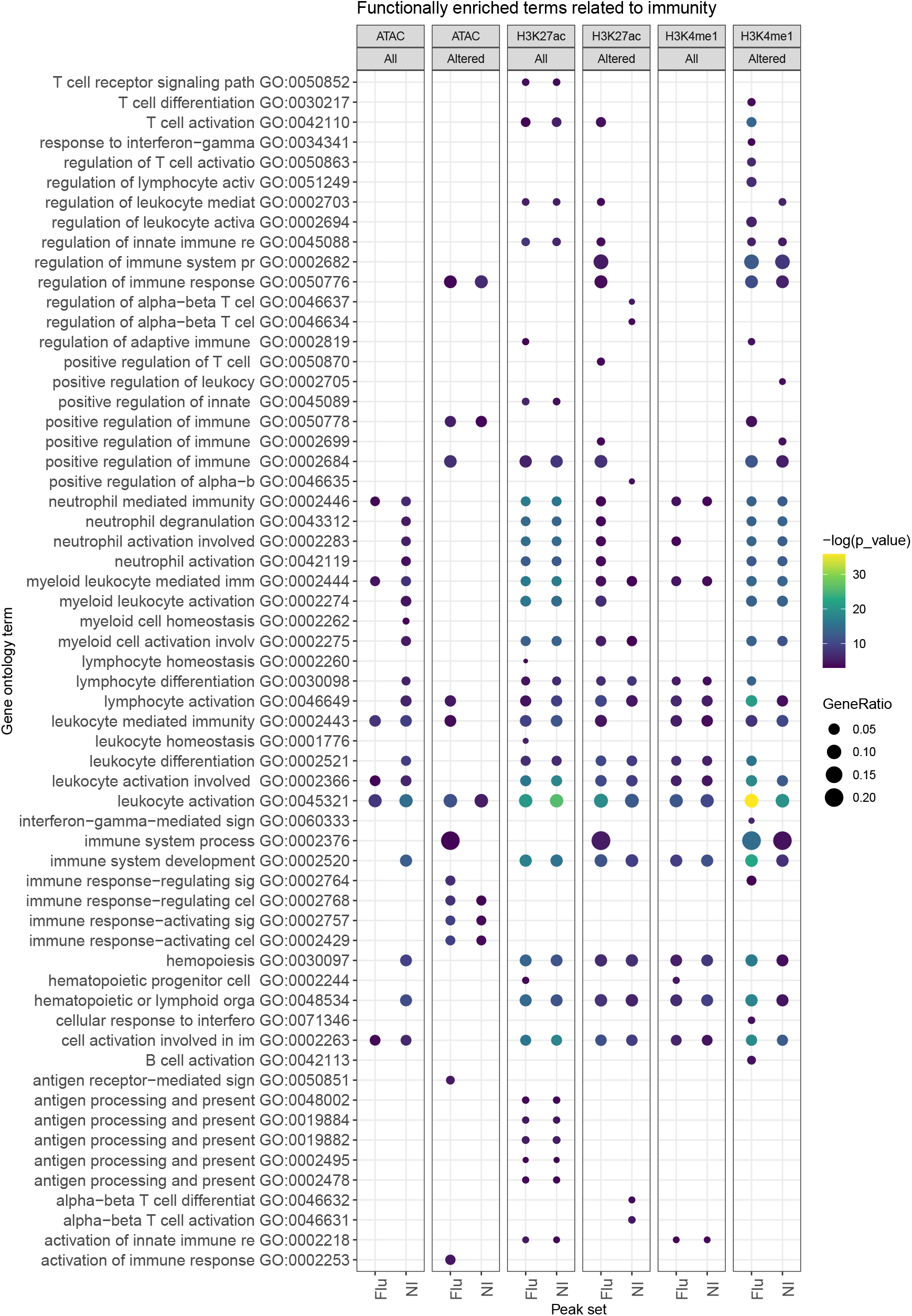
Functional enrichment of genes that are within 10 Kbp of H3K4me1, H3K27ac and ATAC common and altered (graph-only) peaks. We show only the enriched gene ontology terms that relate to immunity and viral infection. The gene ratio is the share of genes that are associated with a GO term.

**Figure S8:**
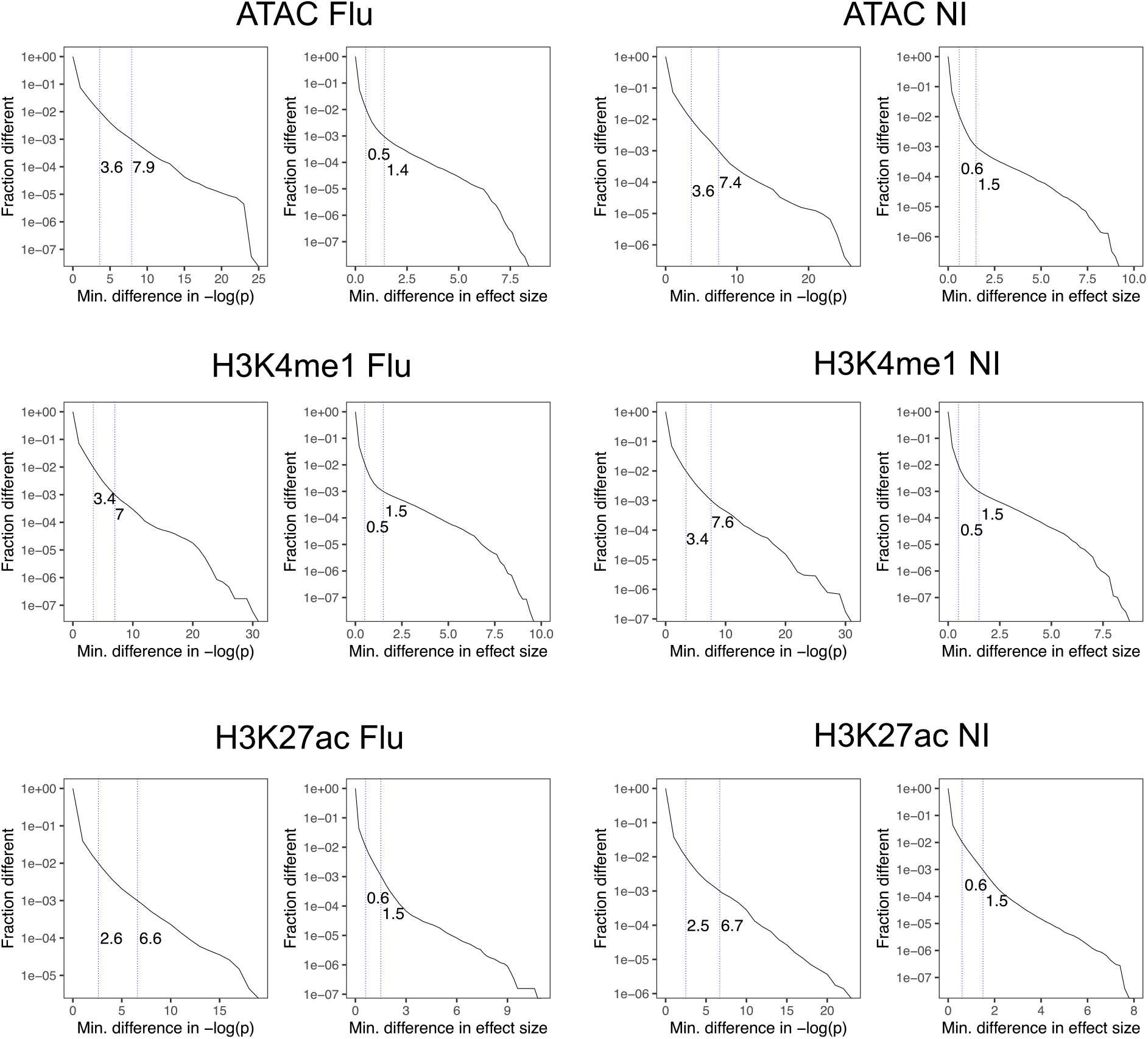
ATAC, H3K4me1, H3K27ac quantitative trait loci are estimated using counts derived from reference and cohort graph alignments in flu-infected and non-infected conditions. We show the fraction of QTLs (y-axis) for which the p-values or effect sizes change by a minimum amount (x-axis). The first line marks the 99th percentile and the second line marks the 99.9th percentile.

**Figure S9:**
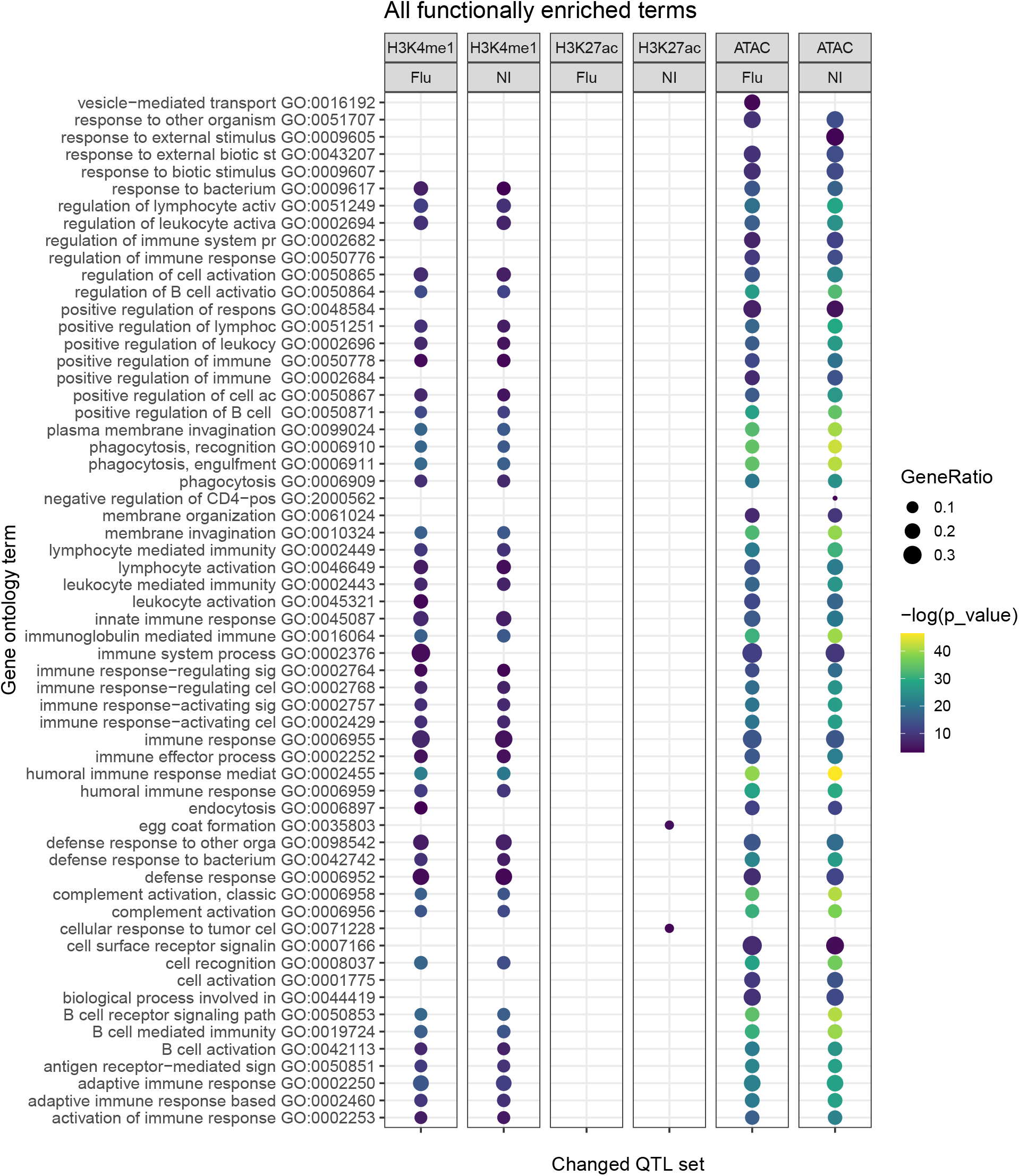
Gene ontology enrichment of genes located within 10 Kbp of peaks associated with hQTLs or caQTLs that change when using a genome graph. Only ca/hQTLs with p-value and effect size changes in the 99.9th percentile were selected. H3K27ac Flu and NI ca/hQTL showed little or no enrichment due a smaller number of ca/hQTLs.

**Figure S10:**
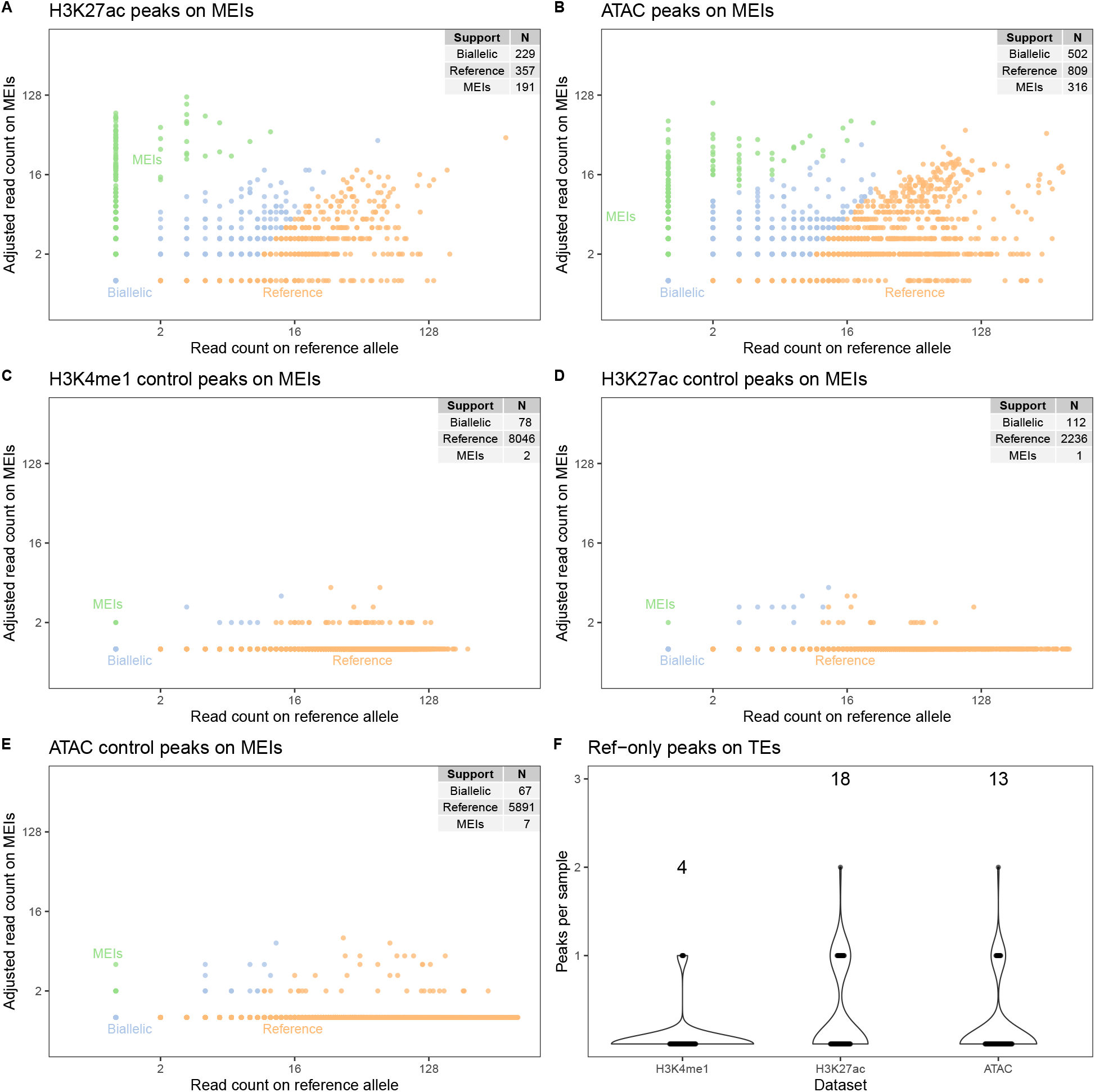
A) B) Partitioning of reads between the reference allele and the alternative allele in peaks that overlap heterozygous or homozygous MEIs in H3K27ac and ATAC-seq. The number of peaks with MEI, biallelic and reference read support is summarized in the table. C) D) E) The same for loci where the samples are known to be homozygous for the reference allele. F) The number of ref-only peaks that overlap MEIs.

**Figure S11:**
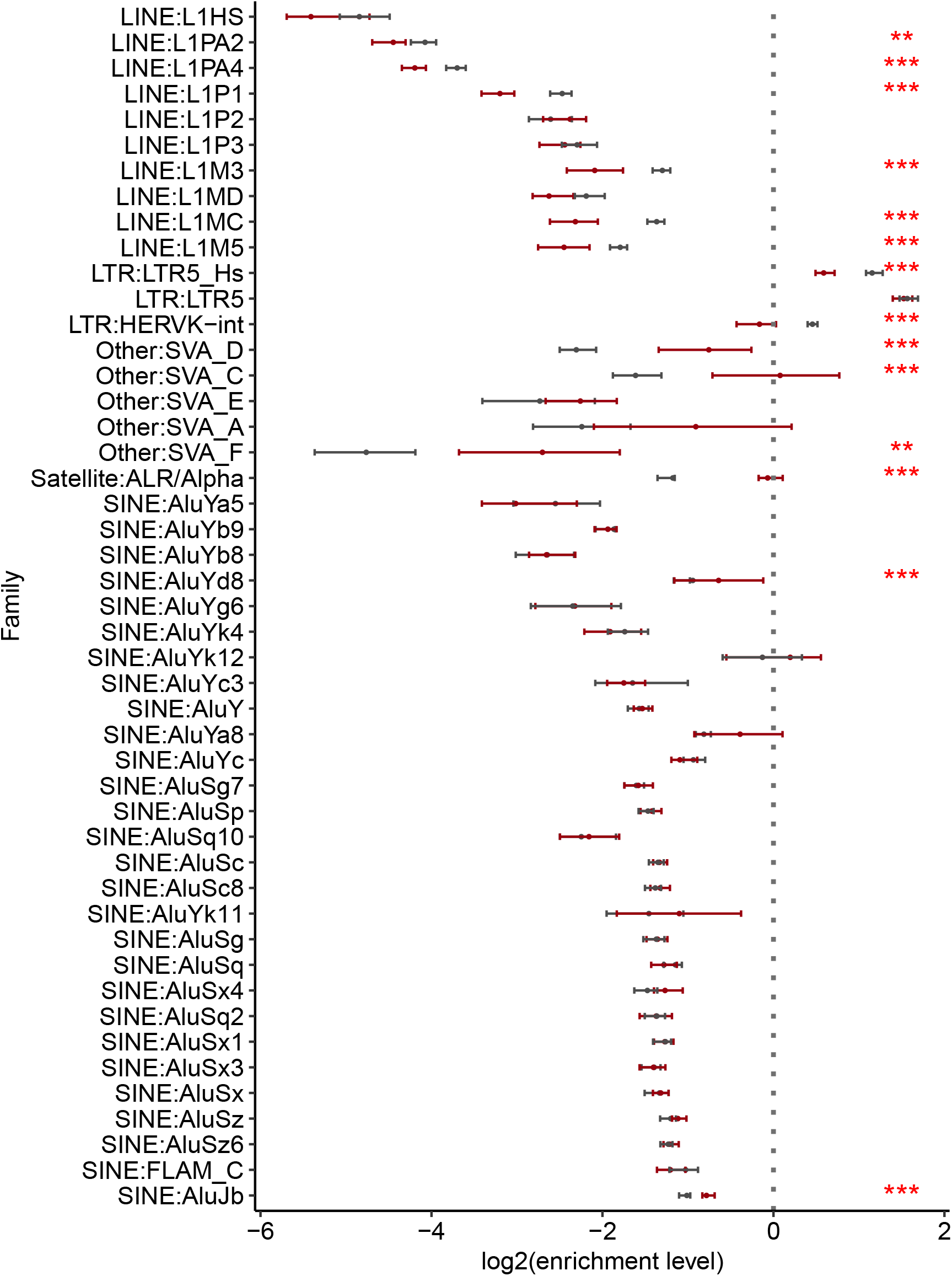
Distribution of enrichment levels of reassembled families within accessible chromatin regions (ATAC) in infected (red) and non-infected (black) samples. SVA families tend become more enriched in accessible chromatin after infection and are highly variable between individuals. Enrichment levels were calculated using methods described by Xun Chen et al [34].

**Figure S12:**
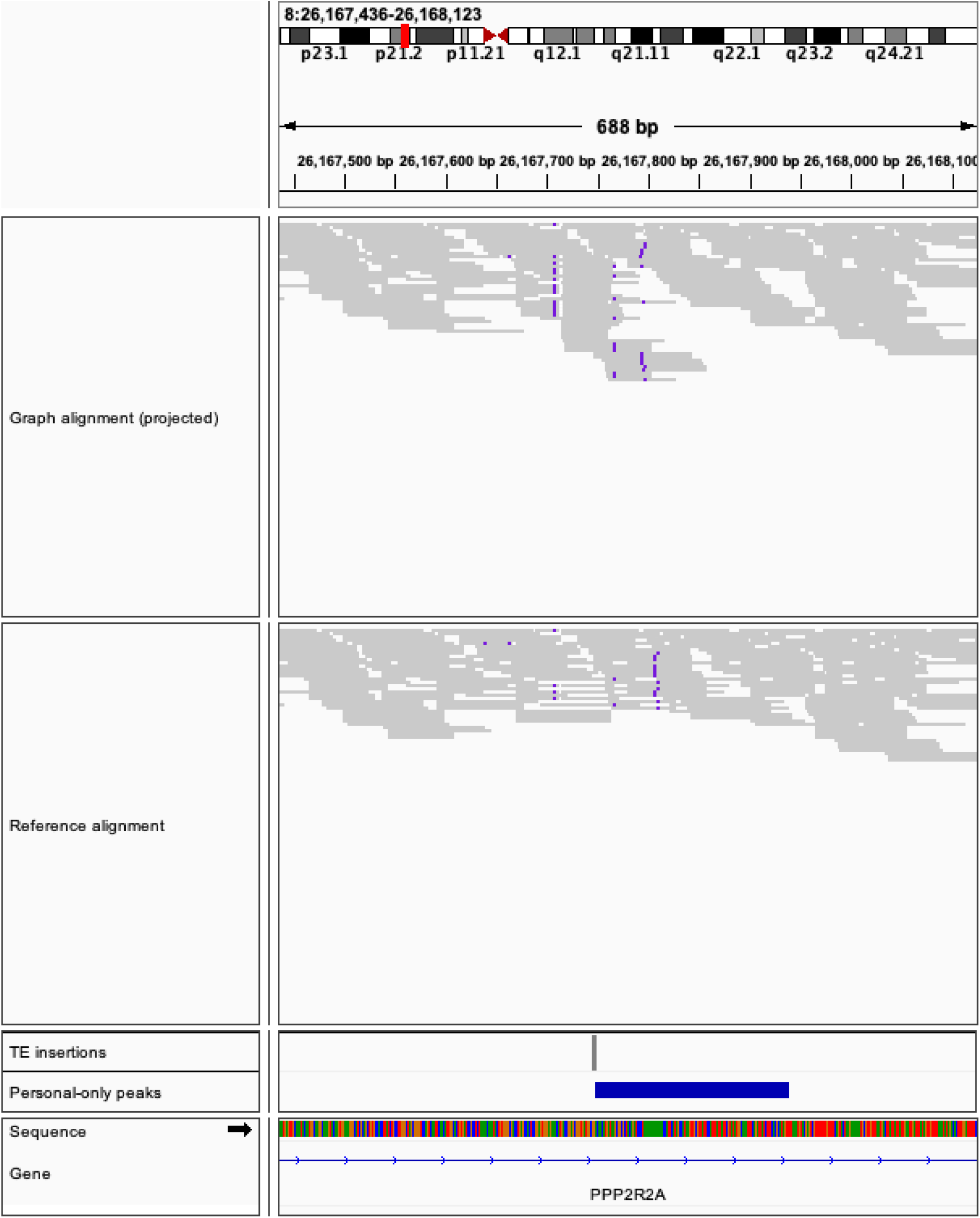
Linear surjection of the example locus (top) versus the reference alignment (bottom) in a Alu polymorphic locus. Note that the actual read pileup is deformed by the surjection to the linear reference. Reads with a MAPQ below 10 have been filtered out. This peak is graph-only in five non-infected samples (AF04, AF08, AF16, EU13, EU47).

**Figure S13:**
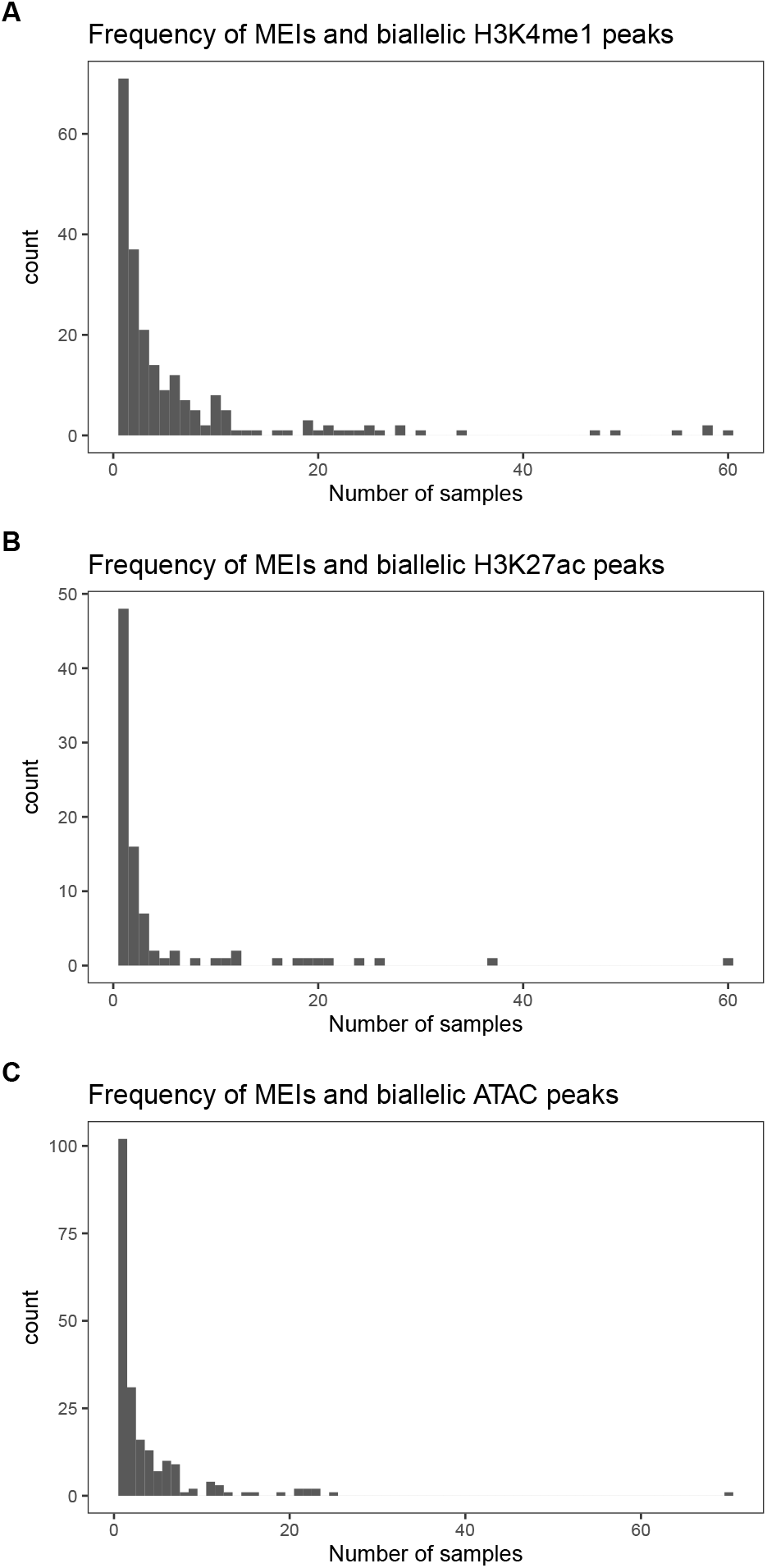
The number times we observe a given MEI and biallelic peak in the cohort at the same locus in A) H3K4me1, B) H3K27ac and C) ATAC-seq.

**Figure S14:**
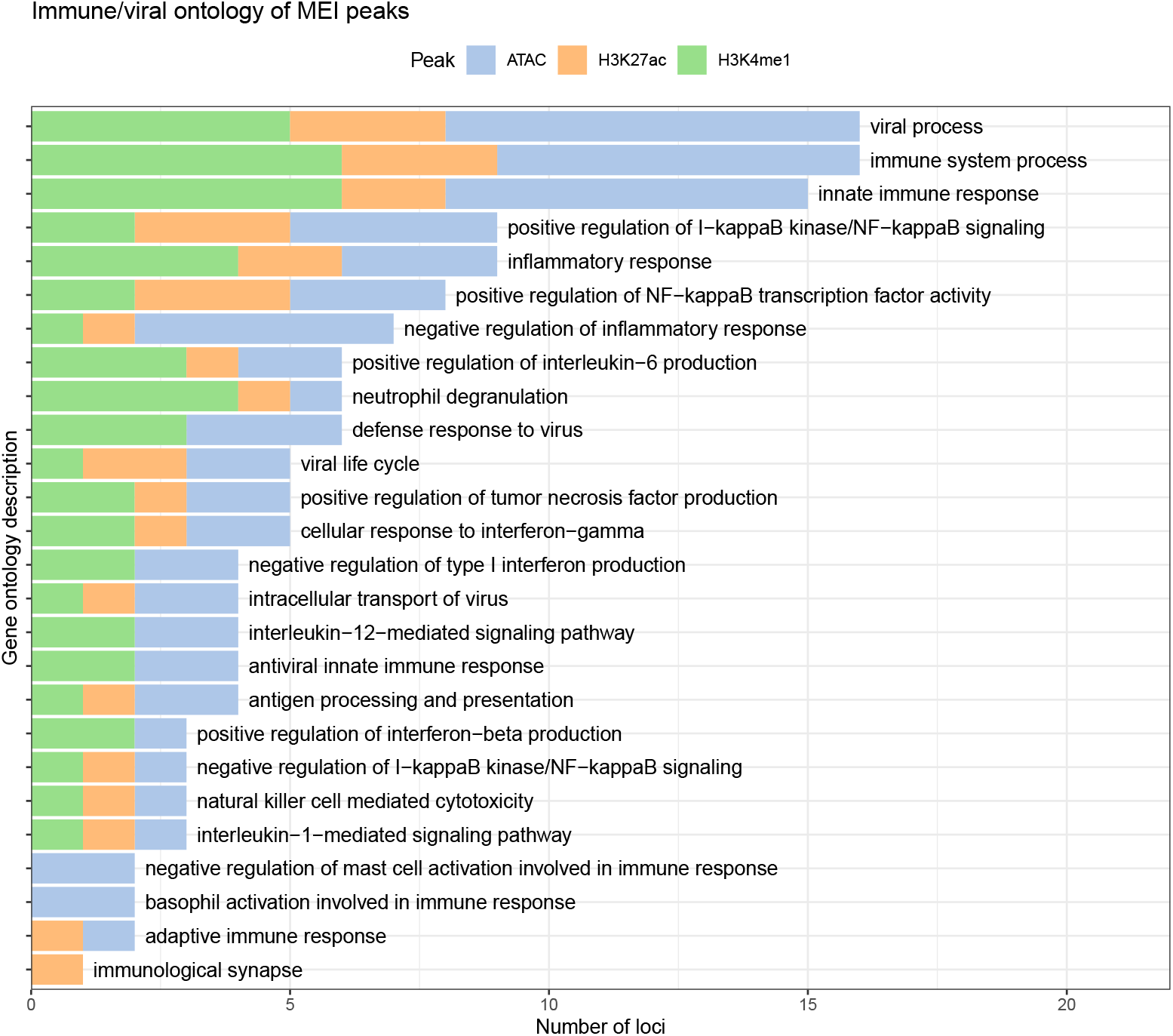
Top ranking GO terms related to immunity and viral infection of genes that are within 10 Kbp of MEI peaks.

**Figure S15:**
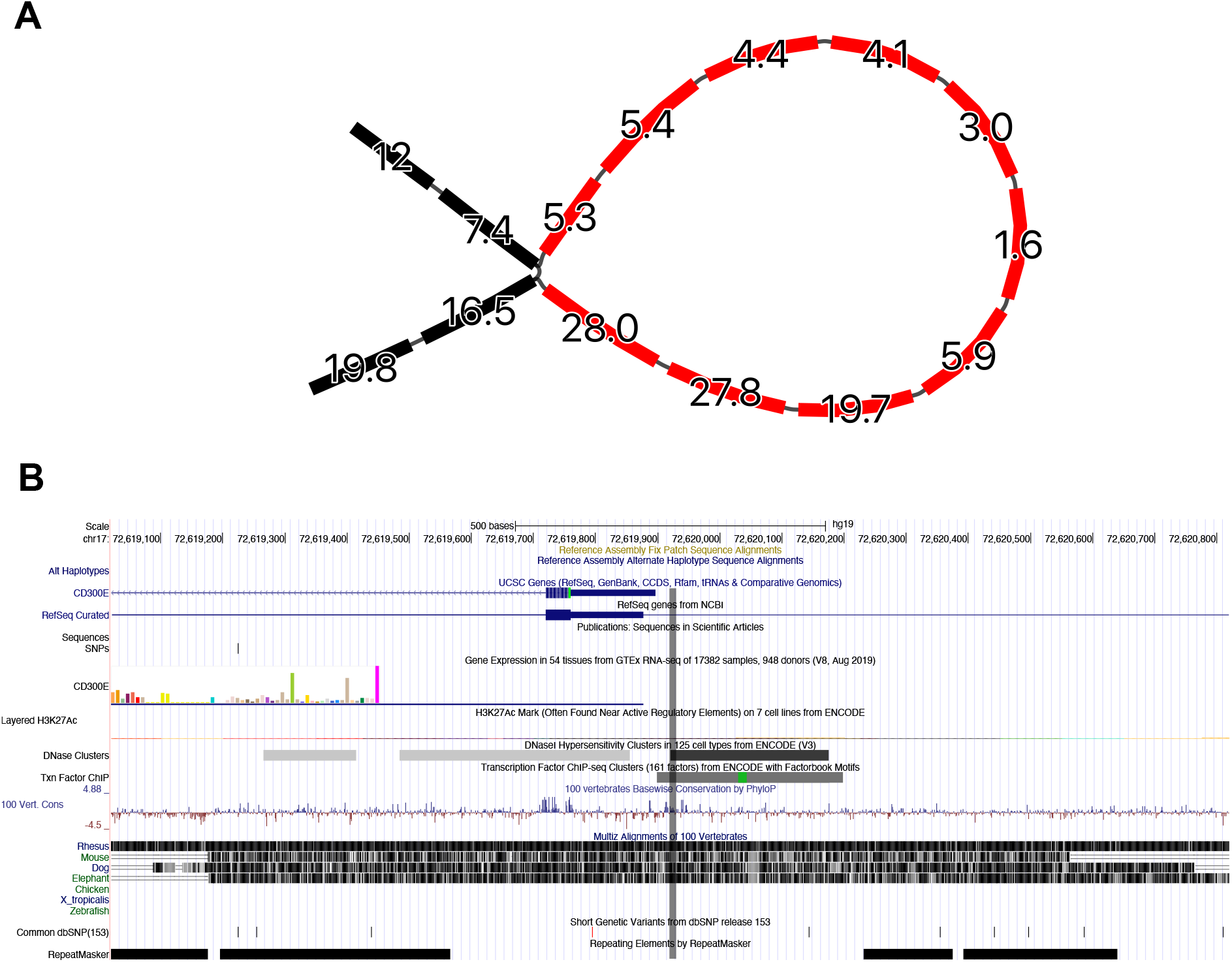
A) Alu mobile element insertion that supports a H3K27ac peak. B) This insertion is immediately upstream of the CD300E gene, and is within a DNase cluster and a ChIP-seq transcription factor cluster. Vertical line marks the site of insertion.

**Figure S16:**
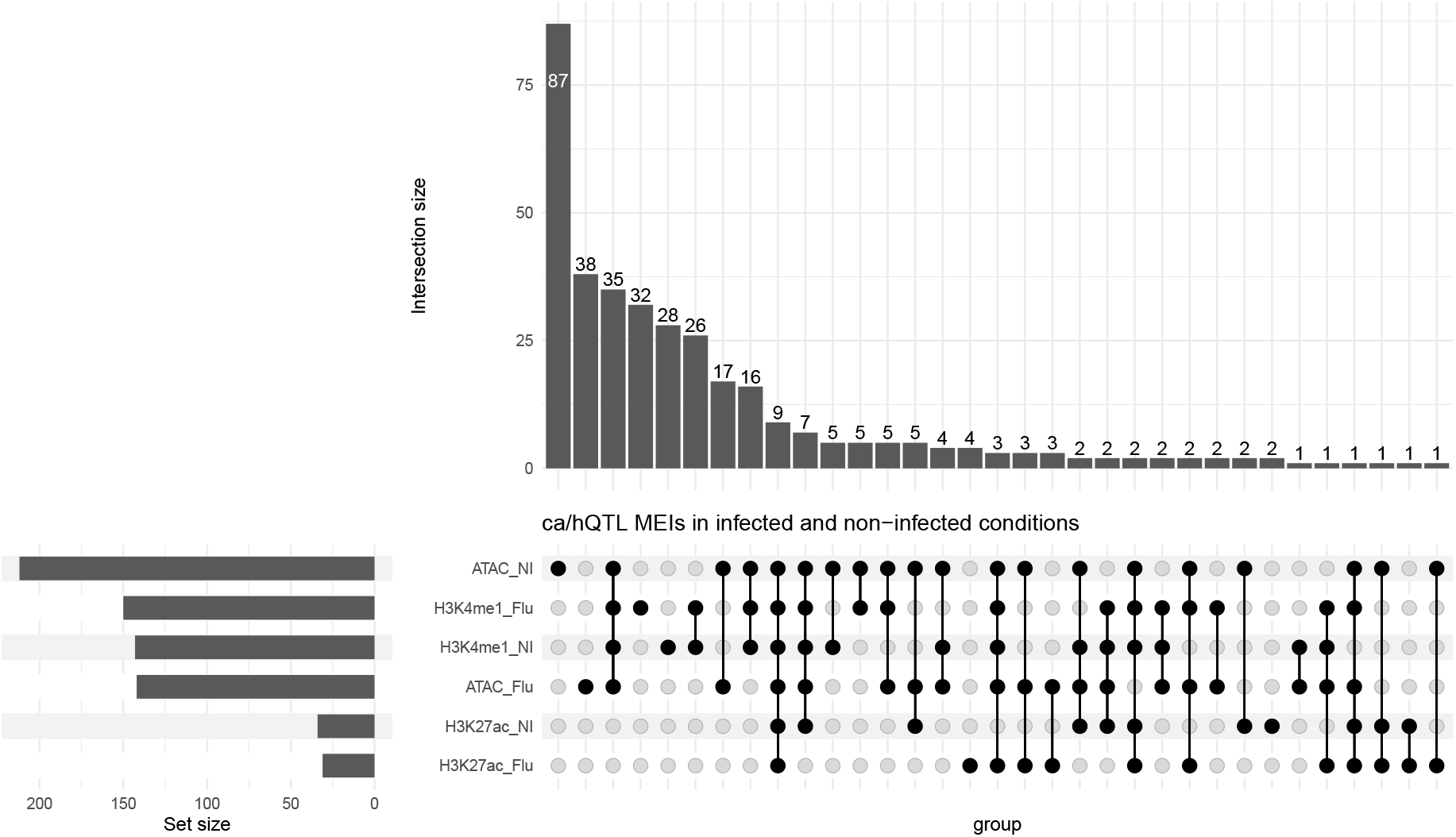
Number of MEIs that are ca/hQTLs in infected and non-infected macrophages.

